# Tetramerisation governs SALL transcription factor function in development and disease

**DOI:** 10.1101/2025.10.27.684836

**Authors:** Sara Giuliani, Kashyap Chhatbar, Martin Wear, Jacky Guy, Tricia Mathieson, Hayden Burdett, Lisa George, Grace Alston, Christos Spanos, Toni McHugh, David Kelly, Raphaël Pantier, Adrian Bird

**Affiliations:** Institute of Cell Biology, University of Edinburgh, Edinburgh, United Kingdom; School of Informatics, University of Edinburgh, Edinburgh, United Kingdom; Edinburgh Protein Production Facility (EPPF), Institute of Quantitative Biology, Biochemistry and Biotechnology, University of Edinburgh, Edinburgh, United Kingdom; Institute of Molecular Plant Sciences, University of Edinburgh, Edinburgh, United Kingdom; Wellcome Discovery Research Platform for Hidden Cell Biology, University of Edinburgh, Edinburgh, United Kingdom; IGBMC, CNRS UMR7104, INSERM U1258, Université de Strasbourg, Illkirch Cedex, France

## Abstract

Spalt-like (SALL) proteins are C2H2 zinc-finger transcription factors important for embryogenesis, with mutations in SALL1 and SALL4 causing rare congenital disorders Townes-Brocks and Okihiro syndromes, respectively. While SALL proteins are known to associate with one another, the biological significance of the resulting complexes is unknown. Here we define a conserved glutamine-rich region that mediates SALL1/4 homo- and heterotetramerisation and find that complex formation is indispensable for DNA binding. Modelling a patient mutation that abolishes SALL4 multimerisation led to gene misregulation and, in mice, embryonic lethality, therefore phenocopying a complete *Sall4* knockout. Furthermore, a common disease-causing SALL1 truncation, which retains multimerisation but lacks DNA-binding domains, sequesters SALL4 into heterotetramers that are defective in DNA binding, thereby providing a mechanistic explanation for the dominant-negative effects of many Townes-Brocks mutations. Together, our findings establish tetramerisation as a prerequisite for SALL function, linking complex formation to developmental gene regulation and human disease.

## Introduction

Spalt proteins were first described in *Drosophila melanogaster* where the two family members, *Spalt-major* and *Spalt-related,* play a role in embryonic patterning^1^. Mammals possess four related Spalt-like proteins, SALL1-4, each featuring several C2H2 zinc-finger clusters (ZFCs) and a N-terminal motif that confers interaction with RBBP4/7 in the Nucleosome Remodelling and Deacetylase (NuRD) co-repressor complex (Figure 1A)^2–5^. SALL1 and SALL4 are expressed in embryonic stem cells (ESCs), required for mouse development and linked to human congenital disorders, while SALL2 and SALL3 are neither expressed in ESCs nor associated with specific diseases^6^. The C-terminal zinc-finger cluster (ZFC4), which is conserved in SALL1, SALL3 and SALL4, mediates DNA binding with specificity for AT-rich motifs (Figure 1A)^7–12^. Disruption of this domain through homozygous mutations in SALL4 abolishes chromatin binding, up-regulates AT-rich genes and leads to precocious differentiation of ESCs. In mice, a homozygous inactivating mutation in ZFC4 causes embryonic lethality, closely resembling *Sall4*-null phenotypes^8,13,14^. Phenotypic defects in animals heterozygous for the *Sall4*-null allele include perinatal lethality^13^ and resemble those seen in Okihiro syndrome (OS), a malformation disorder caused by *SALL4* haploinsufficiency due to heterozygous mutations that generate premature stop codons and loss of functional domains^15–19^. Clinically, OS manifests with upper limb abnormalities, Duane anomaly, renal defects, hearing loss, and congenital heart defects^20^. Notably, only a few heterozygous missense mutations are known to cause OS, all of them located in ZFC4^21,10^, again indicating that interaction with AT-rich DNA is essential for SALL4 function. Less is known about the functional significance of the other ZFCs, although ZFC2 of SALL4 can mediate protein-protein interaction with the BTB/POZ domain-containing protein PLZF^22^. Additionally, thalidomide-driven interaction with the E3 ubiquitin ligase complex is dependent on ZFC1, leading to SALL4 degradation and embryopathies in humans resulting in symptomatic overlap with OS^23,24^.

**Figure 1.**
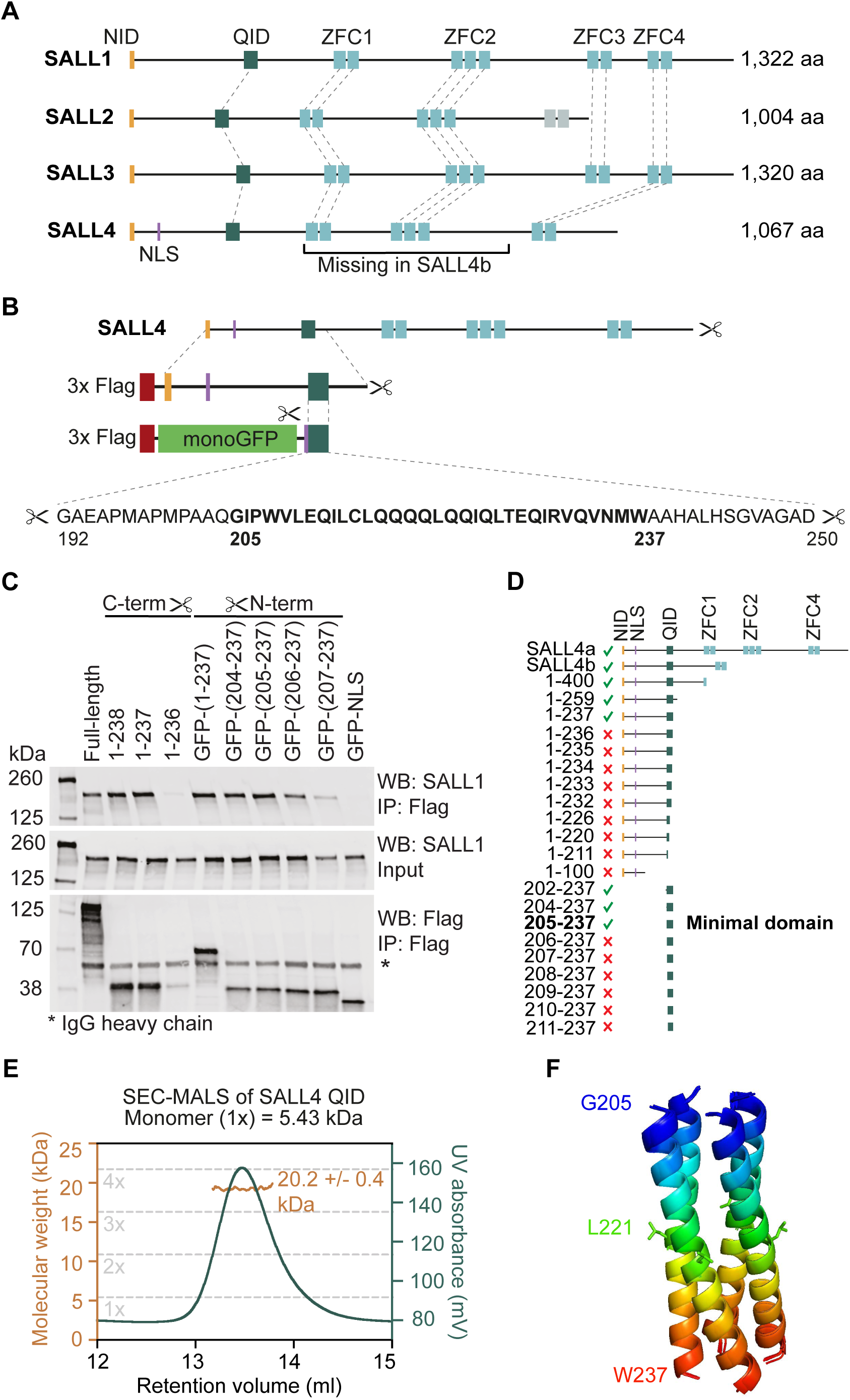
Characterisation of a minimal tetramerisation domain in SALL4. (A) Protein alignment of mouse SALL proteins and their functional domains, including the NuRD-interaction domain (NID), glutamine-rich interaction domain (QID), C2H2 zinc-finger clusters (ZFC1-4) and the nuclear localisation signal (NLS) mapped for SALL4. The domain missing in the SALL4b isoform is bracketed. (B) Cartoon of the experimental design for mapping the QID in SALL4. 3xFlag-tagged SALL4 constructs were used for C-terminal deletions (Figure S1B,C), followed by 3xFlag-GFP-tagged SALL4 construct for N-terminal deletions (Figure S1D). The N-terminal deletions were also fused to the SALL4 NLS to ensure nuclear localisation. The sequence of the QID is shown below, with residue numbers and the minimal domain highlighted. (C) Western blot analysis of co-IPed endogenous SALL1 from transfections of N- and C-terminal deletions as well as Flag-tagged SALL4 full-length into E14/T ESCs. Asterisks indicate IgG heavy chain of the mouse anti-Flag antibody used for IP. Quantified (n=3) in Figure S1E. (D) Diagram of SALL4 full-length (SALL4a) and its functional domains as well as the tested truncation constructs. Ticks indicate that a construct co-IPed endogenous SALL1 and SALL4 while crosses indicate a loss of interaction. The minimal domain construct (205-237) is highlighted in bold. (E) SEC-MALS analysis of recombinant SALL4 QID (192-237) peptide. Green line indicates the UV absorbance at 280nm of the protein elution profile form the SEC while the orange line shows the molecular weight that was measured by the MALS. Grey dashed lines indicate molecular weights expected for monomer (1x), dimer (2x), trimer (3x) and tetramer (4x) assemblies. (F) *In silico* modelling of the SALL4 QID (205-237) tetramer. All 5 models are aligned to each other and coloured from N-to C-term (blue to red). Residue L221 and its sidechain is highlighted in green.

Absence of SALL1 is generally less severe than SALL4-deficiency in mice^13^ and ESCs^14^ but leads to kidney defects^25^. There is however evidence of functional overlap between SALL1 and SALL4, as double mutant ESCs exhibit a more severe loss of pluripotency compared to *Sall4*-null ESCs^14^. Also, the phenotype seen in *Sall4* heterozygous knockout mice is exacerbated in *Sall1/Sall4* double heterozygous mice^13^. In the present context, however, the evidence from genetic analysis of patients with Townes-Brocks syndrome (TBS) is of interest. TBS is caused by heterozygous mutation of *SALL1* creating frameshifts and loss of functional domains and leading to a symptomatic triad of anal, thumb and ear malformations, often combined with renal, genitourinary tract and foot malformations, hearing impairment and congenital heart disorder^26–28^. A *Sall1^ΔZFC1-4^* mouse model mimicking one of the disease-causing mutations^29^ displayed a much stronger phenotype than homozygous *Sall1* knockout (KO) mice^25^ and recapitulated TBS symptoms. These observations led to the proposal that TBS is not caused by SALL1 haploinsufficiency but arises instead through a dominant-negative mechanism^30–32^. It was speculated that this might involve “toxic” interactions with other SALL proteins^29^, especially SALL4 as there is a significant symptomatic overlap between OS and TBS^19,33^.

The present study explores the possibility that the observed genetic interactions are caused by the physical multimerisation of SALL proteins, which is potentially mediated by a glutamine-rich (Q-rich) region near the N-terminus of all SALL proteins (Figure 1A)^29,31^, but whose biological relevance was unknown. We find that the SALL4 interaction domain assembles as a tetramer and that mutations abolishing multimerisation disrupt DNA binding and gene regulation in ESCs as well as embryonic development in mice. Furthermore, we demonstrate that the presence of a disease-causing SALL1 truncation impaired sub-nuclear localisation and gene regulation by SALL4 *in trans*. In summary monomeric SALL proteins are essentially non-functional, as multimerisation is a prerequisite for them to reach the threshold of DNA-binding avidity needed for their physiological activity.

## Results

### Characterisation of a minimal tetramerisation domain in SALL4

To investigate the functional importance of SALL protein multimerisation, we first defined the minimal interaction domain. Previous studies showed that the two isoforms of SALL4 (SALL4a/b) can interact with each other, indicating that the central segment containing ZFC1 and ZFC2 is not required for multimerisation (Figure 1A)^34^. Moreover, a truncation of SALL1 missing all zinc-finger clusters is still able to interact with all SALL proteins in transfection experiments, pointing to their N-terminus as potential interaction domain^29^. In agreement with this conclusion, a study of chicken orthologs of SALL1 and SALL3 found that deletion of 22 amino acids in the conserved Q-rich region abolishes their interaction (Figure S1A)^31^. To precisely map the multimerisation domain, we started with a library of expression vectors carrying 3xFlag-tagged SALL4 with various C-terminal truncations. Following transient transfections into wild-type (WT) ESCs, we tested for co-immunoprecipitation (co-IP) of exogenous SALL4 constructs with endogenous SALL1 and SALL4 by western blot. The interactions were lost after a truncation removing residue 237, a tryptophan downstream of the Q-rich region, thus defining the C-terminal extremity (Figure 1C,D and Figure S1B,C,E). To further refine the mapping of the multimerisation domain, we expressed N-terminal SALL4 truncations (up to W237) fused with a 3xFlag-tagged monomeric GFP^35^ (monoGFP). These experiments defined a minimal 33 amino acid domain essential for interaction with endogenous SALL1/4 (Figure 1C,D and Figure S1D,E) that included the Q-rich domain and also flanking amino acid sequences enriched in hydrophobic residues (Figure S1A). This Q-rich Interaction Domain (QID) is highly conserved between SALL family proteins (SALL1-4) but also across distantly related animal species including Drosophila and human (Figure S1A).

To test whether the isolated QID can self-associate as predicted, we expressed the SALL4 QID fused to a 6xHis-monoGFP solubility tag in *E. coli* (Figure S2A). After cleaving off the tag using TEV protease, a QID-containing peptide (5.3 kDa) was purified by gel filtration (Figure S2B). We determined the stoichiometry of SALL4 complexes using size exclusion chromatography coupled with multi angle light scattering (SEC-MALS), which showed that the QID stably assembles as a tetramer. No other multimeric complexes were detected (Figure 1E). To further investigate the likely mode of interaction, we performed *in silico* modelling using AlphaFold-Multimer^36,37^. The QID is predicted to form an alpha-helix with high confidence and all five models suggest higher order assembly as a four-helical bundle with almost perfect overlap (Figure 1F and Figure S2C,D).

### Multimerisation-deficient SALL4 phenocopies a full SALL4 knockout

To investigate the functional relevance of SALL protein multimerisation, we used ESCs as a cellular model as they show clear phenotypic defects after SALL4 knockout (S4KO), including an unstable stem cell state and global transcriptional deregulation^8,14^. Using CRISPR/Cas9, two different strategies were chosen to disrupt multimerisation of endogenous SALL4 via its QID (Figure 2A and Figure S3A). Firstly, we generated a cell line with homozygous deletion of 22 amino acids within the QID (QIDΔ, Figure 2A)^31^. Additionally, we generated an ESC line carrying a L221P missense mutation (QIDmut, Figure 2A) which is analogous to a human variant identified in an Okihiro syndrome patient (c. 677T>C, p. L226P in *hSALL4,* ClinVar^38^ accession: VCV000423383.2). Contact with clinicians confirmed that the patient had been formally diagnosed with OS and featured classical symptoms such as upper limb malformations including missing thumbs and Duane anomaly. We hypothesised that this proline substitution in the middle of the alpha-helical QID (Figure 1F) would disrupt complex assembly by distorting the helix^39–41^ (Figure S2E). Lastly, we generated a new S4KO ES line on the same genetic background as our other ESCs (Figure S3A), as well as a heterozygous S4KO line (HET S4KO) mimicking haploinsufficiency. We confirmed that mutant SALL4 was still expressed, albeit at slightly reduced levels for QIDΔ and QIDmut cell lines, similar to HET S4KO (Figure S3B). To verify loss of multimerisation, we performed co-IPs with a SALL4 antibody using extracts from our QIDΔ, QIDmut and control lines. As expected, both QIDΔ and QIDmut lost interaction with endogenous SALL1 (Figure 2B). We further verified that the QIDΔ mutation also disrupts homomultimerisation of SALL4 by co-IP following transfection of Flag-tagged and HA-tagged constructs, allowing the distinction between WT and mutant forms (Figure S3C). To test the effect of SALL4 multimerisation mutants on transcription, we performed quantitative RT-qPCR and found that a selection of SALL4 target genes^8^ are deregulated in QIDΔ and QIDmut lines, but this effect was not seen in HET S4KO ESCs, which express ∼50% WT levels of SALL4 (Figure S3D). For a comprehensive analysis of the mutant transcriptomes, we performed RNA-seq. Strikingly, this showed that a large proportion of deregulated genes are shared between QIDΔ, QIDmut and S4KO (Figure 2C), with the magnitude of transcriptional deregulation (log_2_ fold change versus WT control) showing strong correlations across all genotypes (Figure 2D and Figure S3E,F). Importantly, this indicates that mutational inactivation of the SALL4 QID leads to an almost complete loss of function of this transcription factor (TF).

**Figure 2.**
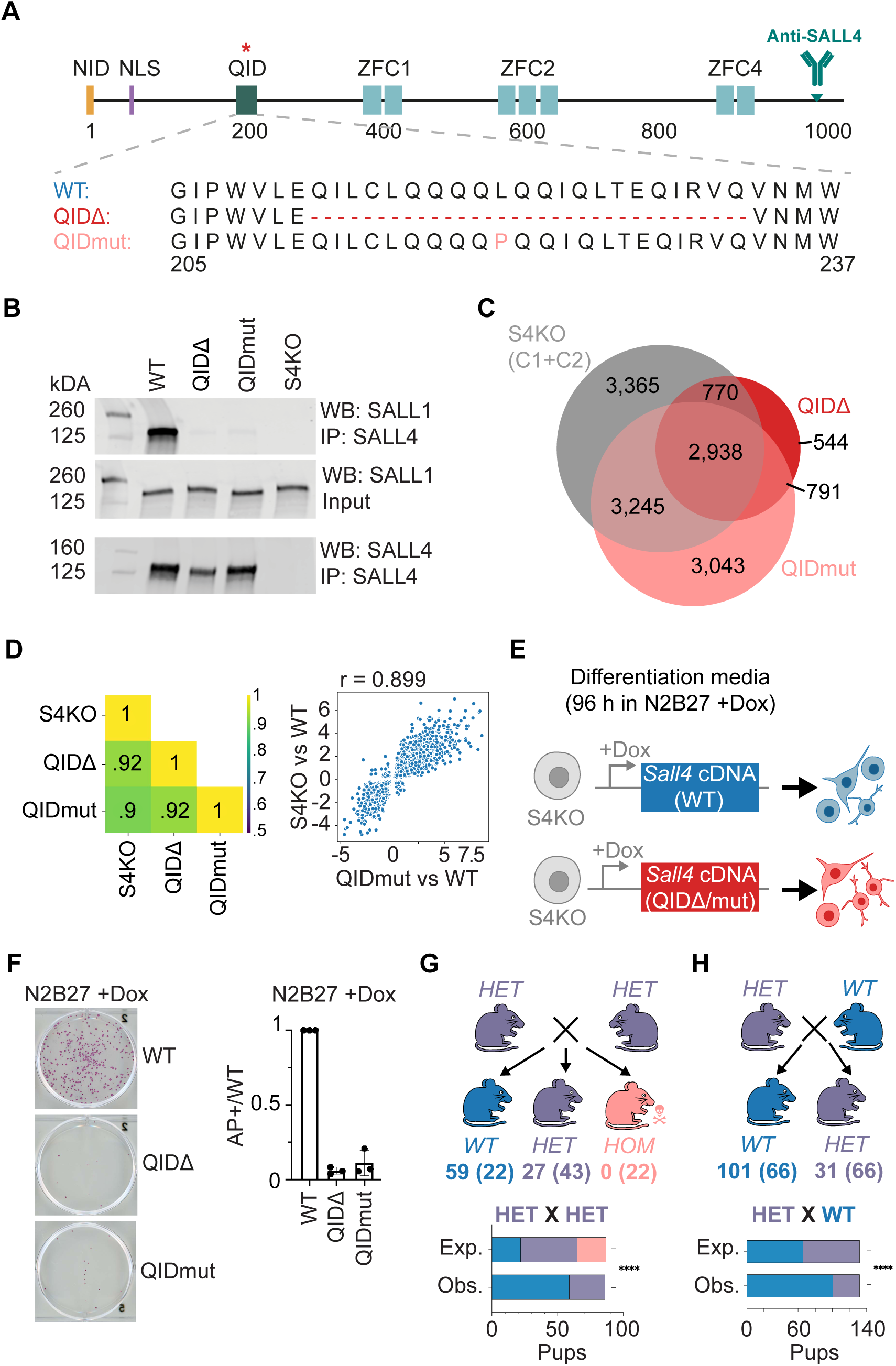
Multimerisation-deficient SALL4 phenocopies a full SALL4 knockout. (A) Diagram of SALL4 with its functional domains and the antibody epitope. Mutations designed to disrupt the QID are highlighted in red. The CRISPR strategy used to generate the cell lines is detailed in Figure S3A. (B) Western blot analysis of co-IPed SALL1 from a SALL4 IP with the polyclonal rabbit anti-SALL4 antibody in QIDΔ, QIDmut and control cell lines. (C) Venn diagram showing the overlap of genes that are deregulated in S4KO (clones C1+C2), QIDΔ and QIDmut cell lines compared to WT ESCs (padj>0.05). (D) Plot showing the Pearson correlation values (r) of the 2,938 shared deregulated genes between all mutant lines. One dot plot (with one dot representing the log_2_ fold change of one gene vs WT in both genotypes) of the S4KO – QIDmut comparison is shown illustrating that many genes are deregulated in the same way in both genotypes (for more details see Figure S3E,F). (E) Workflow of the spontaneous differentiation experiment using piggyBac cell lines, where Sall4 WT (blue) and QIDΔ and QIDmut (red) cDNAs were integrated into S4KO ESCs. Addition of doxycycline (+Dox) to the culture medium switches transgene expression on, leading to overexpression of SALL4. (F) Representative wells of stem cell colonies stained by alkaline phosphatase staining^44^ of all the different piggyBac cell lines after 96 h in N2B27+Dox medium. The quantification on the right shows the mean of the number of AP-positive colonies (AP+) found in each genotype and normalised to WT (positive control) for three biological replicates (individual data points). The error bars indicate the standard deviation (SD). (G) Diagram showing the results of genotyping from pups (P21) following crosses of heterozygous (HET) QIDmut animals. No homozygous (HOM) animals were recovered, indicating complete embryonic lethality. The bar graph below shows quantification and statistical comparison with expected Mendelian distribution using the Fisher’s exact test (p-value< 0.0001, ****). H) Diagram and quantification of WT by HET crosses is shown illustrating the QIDmut heterozygous animals are also born at a sub-Mendelian ratio (Fisher’s exact test, **** = p-value < 0.0001).

As the overexpression of WT SALL4 decreases spontaneous differentiation of ESCs in N2B27^14^ and is also known to reprogram cells to pluripotency^42^, we decided to test the importance of the QID during this cellular process. Using the piggyBac system^43^, we randomly integrated inducible constructs carrying either WT, QIDΔ or QIDmut *Sall4* cDNA into S4KO ESCs^14^ (Figure 2E) and established that the addition of doxycycline (+Dox) to the medium led to SALL4 overexpression (Figure S3G). We then cultured a fixed number of ESCs for each genotype in N2B27 differentiation medium^14,8^ +Dox for 96h, during which the SALL4 version was overexpressed. Colonies were stained for alkaline phosphatase (AP) expression, a marker of pluripotent cells^44^. Consistent with previous findings^14^, WT SALL4 overexpression efficiently reduced ESC differentiation, as demonstrated by the large number of AP-positive colonies (Figure 2F). In contrast, QIDΔ or QIDmut SALL4 did not reproduce this phenotype (Figure 2F and Figure S3H), again highlighting the loss of function of monomeric SALL4.

To further test the role of SALL4 tetramerisation *in vivo*, we generated a mouse line carrying the L221P missense mutation (QIDmut). Intercrossing of heterozygous animals (HET x HET) demonstrated that this mutation is lethal in the homozygous state, as no pups with this genotype were born (Figure 2G). We further noticed that heterozygous animals were born at sub-Mendelian ratios, phenocopying effects already seen in S4KO animal models^13^ (Figure 2H). Taken together, these phenotypic assays show that QID mutant SALL4 phenocopies a full S4KO and therefore appears to be non-functional. Of further interest, the L221P mouse mimicking the human patient mutation c. 677T>C represents the first animal model for an OS-causing missense mutation. This atypical mutation has previously been classified as a “variant of uncertain significance” and our results therefore provide a molecular explanation for the disease phenotype observed in this patient. Of note, the only other known missense mutations causing OS are clustered within the DNA-binding domain ZFC4^10,21^.

### Monomeric SALL4 cannot bind to DNA efficiently

We next sought to determine the molecular basis of the observed requirement for multimerisation and tested whether SALL4 tetramerisation increases its DNA-binding affinity. It is known that SALL4 co-localises with nuclear foci corresponding to chromocenters in mouse cells, due to its affinity for AT-rich satellite DNA repeats^7,13^. This co-localisation is lost when the SALL4 DNA-binding domain ZFC4 is mutated (ZFC4mut), reflecting a general loss of DNA binding genome-wide^8^. Strikingly, we found that both QIDΔ and QIDmut were unable to localise to heterochromatic foci, indicating a loss of DNA-binding affinity despite the presence of a functional ZFC4 domain in both mutant proteins. This effect is unlikely to be explained by reduced SALL4 protein levels in the mutant lines, as our HET S4KO ESC control (≈50% WT SALL4 expression levels) showed a normal localisation of SALL4 to chromocenters (Figure S3B). To directly assess SALL4 chromatin binding we performed CUT&RUN experiments^45^, which showed a total loss of binding to DNA for both SALL4 QIDΔ and QIDmut, similar to the S4KO and IgG negative controls. This dramatic loss of DNA binding was observed whether the analysis was focused on detected peaks (Figure 3C and Figure S4A) or dispersed genomic binding to AT-rich DNA (Figure S4B). As a control, HET S4KO cells confirmed that even with reduced protein levels SALL4 retains binding to WT peaks, albeit with the expected 50% loss in signal (Figure 3C and Figure S4A), and also retains its affinity for AT-rich regions genome-wide (Figure S4B). Together, these results demonstrate that the SALL4 QID is essential for chromatin binding *in vivo*. A likely explanation is that the DNA-binding affinity of a single SALL4 molecule is below the threshold required for functional chromatin binding *in vivo*, whereas tetramerisation increases DNA-binding avidity via multivalent interactions^46^.

**Figure 3.**
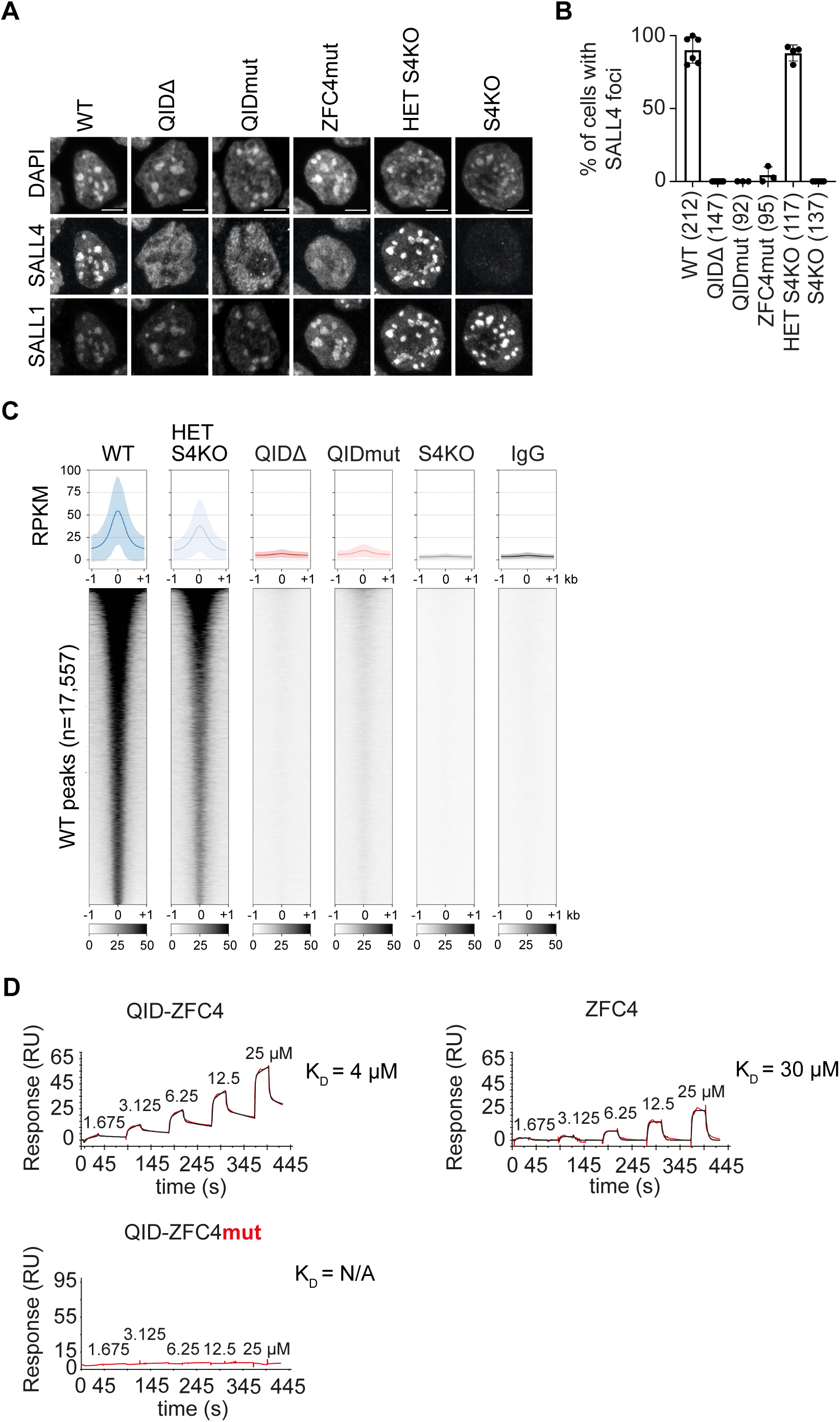
Monomeric SALL4 cannot bind to DNA efficiently. (A) Representative nuclei of the different genetically modified ES cell lines after co-immunofluorescence of SALL4 and SALL1. DNA was stained with DAPI. Scale bar = 5 µm. (B) Quantification of all immunofluorescence microscopy experiments, as shown in panel A. All cells were assessed for having SALL4-positive foci or not and results were expressed as ratios over the total cell numbers (n, shown in brackets next to each genotype). Individual data points indicate independent replicate experiment with error bars representing the SD. (C) Heatmaps and average plots of SALL4 CUT&RUN signal over peak regions (17,557 peaks observed in WT cells) in ESC lines from different genotypes. The Reads Per Kilobase per Million mapped reads (RPKM) are all normalised to a *E. coli* DNA spike-in that was added to each sample. The shaded area in the plots corresponds to the SD of peak signal across all analysed loci. (D) SPR analysis of several SALL4 QID-ZFC4 fusion proteins binding to a DNA probe containing a single ATATTA motif. Mutations in functional domains are highlighted in red and for each construct a K_D_ calculated from a kinetic model is given, unless no binding was detected (N/A).

To directly test this hypothesis, we assessed the *in vitro* DNA-binding potential of minimal constructs including SALL4 QID fused via a flexible glycine linker to the ZFC4 DNA-binding domain, in WT or mutated states (Figure S4C,F). The stoichiometry of each construct was investigated by SEC-MALS, confirming the expected assembly of “QID-ZFC4” and “QID-ZFC4mut” as tetramers, while “QIDmut-ZFC4” and “ZFC4” remained monomeric (Figure S4D). Fusion proteins were then assayed by surface plasmon resonance (SPR) to determine their DNA-binding potential using a probe containing a single ‘ATATTA’ motif with high chip DNA density, likely permitting multivalent interactions. Qualitatively, QID-ZFC4 bound DNA most strongly, followed by ZFC4 alone (Figure 3D). As expected, QID-ZFC4mut showed no binding, confirming that the QID is not a DNA-binding domain. Binding was also reduced for QIDmut-ZFC4, but non-specific interactions with the chip surface - potentially due to exposed hydrophobic residues in the monomeric QID - complicated interpretation (Figure S4E). Overall, the data suggests that tetramerisation of the SALL4 DNA-binding domain ZFC4 increases affinity to DNA compared to monomeric SALL4.

### Binding to the NuRD co-repressor complex is reduced in QID mutant SALL4

In addition to studying the effects of QID mutation on SALL4 DNA-binding activity, we tested its impact on protein-protein interactions. As subunits of the NuRD co-repressor complex are among the strongest SALL4 interactors in ESCs^14^, we first checked by co-IP and western blot analysis whether its core subunit MTA1 interacts with SALL4 QIDΔ and QIDmut and found that this interaction is retained, but reduced (Figure 4A,B). Mass spectrometry analysis of SALL4 immunoprecipitated nuclear protein extracts in WT and mutant cell lines (Figure 4C-E) quantified this reduction as a ∼4-fold decrease in affinity between SALL4 and all known subunits of the NuRD co-repressor complex in both SALL4 QIDΔ and QIDmut cell lines (Figure 4G). In this experiment we verified the complete loss of multimerisation with other SALL proteins after mutation of the SALL4 QID (Figure 4F). From this analysis, we conclude that tetramerised SALL4 is more efficient at recruiting the NuRD co-repressor complex compared to its monomeric form, yet this loss of interaction is less pronounced than the dramatic loss of genome binding.

**Figure 4.**
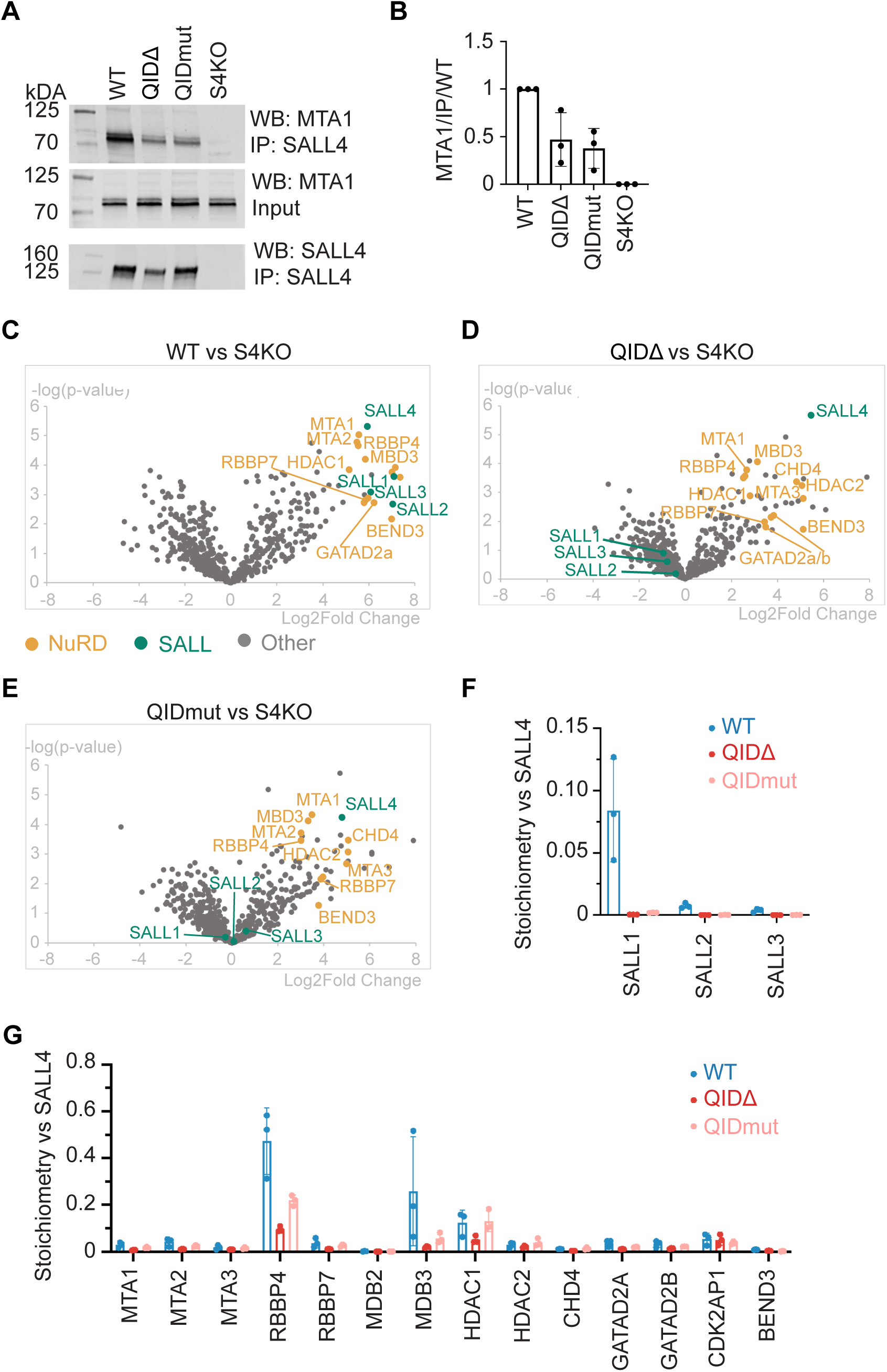
Binding to the NuRD co-repressor complex is reduced in QID mutant SALL4. (A) Western blot analysis of co-IPed MTA1 from a SALL4 IPs with the polyclonal rabbit anti-SALL4 antibody in QIDΔ /mut and control cell lines. (B) Quantification of the co-IP of MTA1 with SALL4 WT or QIDΔ /mut protein, as shown in panel A. The data was acquired in three independent replicates (data points), of which the mean is shown. Error bars represent the SD. (C, D, E) Volcano plot showing all proteins detected by mass spectrometry analysis in SALL4 IPs from WT (C), SALL4 QIDΔ (D) and SALL4 QIDmut (E) ESC lines. Relative enrichment (Log2 Fold Change) and statistical significance (-log(p-value)) were calculated in comparison to S4KO negative control samples. SALL proteins are highlighted in green and NuRD co-repressor complex subunits in yellow. (F, G) Stoichiometries of SALL proteins (F) and NuRD co-repressor complex subunits (G) in SALL4 IPs from WT (blue), QIDΔ (red) and QIDmut (salmon) cell lines, as assessed by mass spectrometry analysis. For each protein, label-free absolute quantification (iBAQ values) was obtained and normalised to the corresponding IPed SALL4 levels. Experiments were performed in technical triplicates (individual data points) and error bars show the SD.

### SALL protein tetramers defective in DNA binding can explain the dominant-negative effect of SALL1 truncations underlying Townes-Brocks syndrome

TBS is caused by mutations that lead to premature stop codons in the SALL1 coding sequence. Such mutations appear to cluster within a “hotspot”, potentially resulting in the expression of a truncated form retaining the multimerisation domain (i.e. QID), but excluding all ZFCs^47^. Notably, a recent study showed that patients with deletions of the entire coding sequence of *SALL1* have milder symptoms than those caused by hotspot mutations^32^, further supporting the argument that these SALL1 protein truncations may exert a dominant-negative effect (see also Introduction). It was speculated that the QID region might underlie the dominance of TBS mutations through an unknown mechanism^29–31^.

We first revisited the evidence for a mutational hotspot by collating recent data regarding TBS-causing mutations from published literature and the ClinVar database (Table S1). In agreement with previous literature^47^, 61 out of 95 *SALL1* mutations (64%) caused a frameshift within this hotspot region (Figure 5A). This mutational pattern is particularly impressive given that the hotspot region, starting immediately downstream of the QID, only includes 20% of the coding sequence. In addition to this, we found that *SALL1* is depleted of similar mutations in the healthy population, as a gnomAD^48^ database (v4.1.0, https://gnomad.broadinstitute.org/) search identified only three variants, all at the C-terminus of the protein after SALL1 ZFC4 (Figure 5A highlighted in blue, namely p. Arg1192* with five cases, p. Leu1197fs6* with 15 cases and p.Met1203* with three cases).

**Figure 5.**
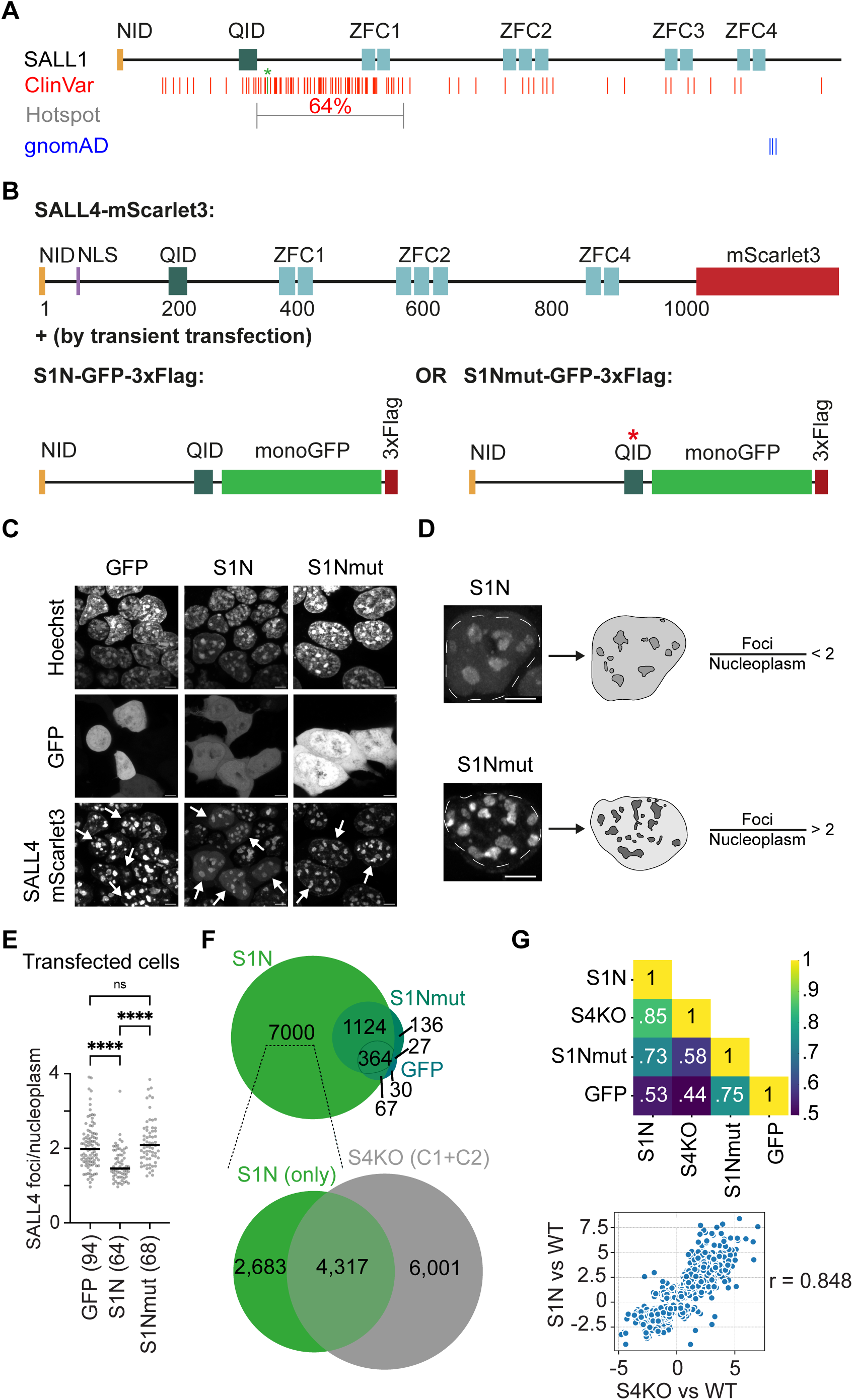
A TBS-causing SALL1 truncation impairs SALL4 sub-nuclear localisation and gene regulation. (A) Diagram of SALL1 with its functional domains as well as the location of TBS-causing non-sense and frameshift mutations in red (Table S1) and neutral mutations found in the general population (gnomAD^48^) in blue. Of note, missense mutations are not shown in this diagram. The modelled 826C>T mutation (S1N) is highlighted in green and with a star. Lines indicate the first mutated residue with missense tails not being shown. Below the disease-causing variants, a grey brackets indicates the location of the previously defined hotspot region^47^. (B) Depiction of SALL4-mScarlet3 fusion protein expressed by genetically engineered ESCs and the human SALL1 constructs that were transiently transfected with C-terminal GFP 3x-Flag tags. The mutation introduced within SALL1 QID (I241P, homologous to mouse SALL4 L221P) disrupts homo- and heteromultimerisation with SALL proteins (see Figure S5C). (C) Representative pictures of live cells expressing endogenously SALL4-mScarlet3 cells and transfected with GFP-tagged constructs: GFP, S1N or S1Nmut All samples were analysed with the same brightness/contrast indicating different transfection efficiencies. In the SALL4 mScarlet3 channel, white arrows highlight transfected cells. (D) Zoom on single nuclei, either transfected with S1N or S1Nmut. The cartoon in the middle depicts how the automated segmentation and quantification programme measured area-normalised SALL4 signal in foci and nucleoplasm. Relative enrichment was quantified as a fluorescence ratio, as shown on the right. (E) Quantification of all live-cell imaging data from (n=5) replicate experiments. Only transfected cells were analysed with each data point representing SALL4 fluorescence ratio (see panel D) in a single cell and the black line representing the median. Statistical analysis was performed using an ordinary one-way ANOVA test with a p-value lower than 0.05 being considered statistically significant (**** p< 0.0001). (F) Venn diagram showing the overlap of genes that are deregulated WT cells transfected with S1N, S1Nmut or GFP. The 7,000 genes that are only deregulated in S1N (not GFP or S1Nmut) are then overlapped with genes deregulated in S4KO (clones C1+C2). All deregulated genes are obtained from comparisons to WT ESCs (padj>0.05). (G) Plot showing the Pearson correlation values (r) of the 4,317 shared deregulated genes between all cell lines. Additionally, the dot plot (with one dot representing the log_2_ fold change of one gene vs WT in both genotypes) of the S1N – S4KO comparison is shown illustrating that many genes are deregulated in the same way (for more details see Figure S5F).

As most pathogenic SALL1 truncations retain the tetramerisation domain but lack DNA-binding ZFCs, we hypothesised that they may lead to the formation of SALL protein tetramers that are defective in DNA binding. A key prediction is that SALL1 truncations would interfere with the DNA-binding capacity of non-mutated SALL1/2/3/4, which are its primary multimerisation partner. As ESCs predominantly express SALL4 (Figure S5A), we decided to express a pathogenic SALL1 truncation in this system to assess the consequences of forced SALL1:SALL4 heterotetramerisation. We selected the most prevalent TBS-causing mutation (c. 826C>T) which introduces a nonsense mutation at arginine 276 in human *SALL1* (Figure 5A, highlighted in green*), leading to the expression of an N-terminal SALL1 fragment^30^ (S1N). As a negative control, we generated a QID-mutant version of this construct (S1Nmut, I241P) and verified by co-IP that it was unable to multimerise with SALL4 (Figure S5C). First, we assessed SALL4 and SALL1 co-localisation with AT-rich heterochromatic foci by immunofluorescence. This showed that SALL4 nuclear signal in S1N transfected cells was entirely diffuse (Figure S5D), suggesting a significant DNA-binding defect of endogenous SALL4 in the presence of the TBS-causing mutation. To validate this observation in live cells, we designed a system that allowed for imaging of both endogenous SALL4 and transfected SALL1 truncations using fluorescent tags (Figure 5B and Materials and Methods). This analysis showed that in cells transfected with the SALL1 truncation S1N, SALL4 was apparent within DNA foci but the signal appeared more diffuse compared to the GFP and S1Nmut controls (Figure 5C). We assessed this phenomenon quantitatively by performing automated image segmentation and calculating the ratio of SALL4 fluorescence signal in foci versus nucleoplasm in each experimental condition (Figure 5D). SALL4 enrichment within foci was significantly reduced in cells transfected with S1N with a median ratio ≈1.5 (versus ≈2 for GFP only control) (Figure 5E). In contrast, SALL4 distribution in cells transfected with S1Nmut was comparable to the GFP control with a median ratio of ≈2.1 (Figure 5E), and there was also no detectable difference in untransfected cells (Figure S5B). The results in both live and fixed cells strongly support the hypothesis that TBS-causing SALL1 truncations reduce the affinity of SALL4 for DNA via QID-mediated heteromultimerisation.

To investigate transcriptional consequences of SALL1 TBS truncation, we used the same experimental approach, but specifically enriched the population of transiently transfected cells by puromycin selection. Initial analysis by RT-qPCR showed that two SALL4 target genes (identified in previous RNA-seq datasets^8^), *Lix1* and *Sox6*, were upregulated in cells transfected with S1N, similarly to S4KO ESCs (Figure S5E). This upregulation was not observed for cells transfected with GFP or S1Nmut controls and was also less pronounced for HET S4KO (expressing ≈50% SALL4 levels). To extend our analyses to the entire transcriptome, we performed RNA-seq, revealing extensive gene deregulation in ESCs transfected with S1N, which was much less pronounced in GFP or S1Nmut controls (Figure 5F). More importantly, a majority of deregulated genes were shared between S1N overexpressing cells and S4KO ESCs, with highly correlated levels of transcriptional deregulation (Figure 5F,G and Figure S5F). This strongly indicates that TBS-causing SALL1 truncation exerts a dominant-negative effect on endogenous SALL4. This effect is dependent on a functional QID, and therefore SALL1:SALL4 multimerisation, as similar correlations were not detected for S1Nmut (Figure 5F,G and Figure S5F). In conclusion, our investigations have now clarified the molecular mechanism underlying the dominant-negative effect of TBS-causing mutations. The expression of pathogenic SALL1 truncations is likely to interfere with non-mutated SALL4 via heteromultimerisation involving a functional QID domain, leading to loss of DNA binding and transcriptional deregulation of SALL4 target genes. Overall, we propose that tetramerisation is a fundamental requirement for high-affinity DNA binding of SALL proteins, and that its disruption can be the root of human congenital disorders OS and TBS (Figure 6).

**Figure 6.**
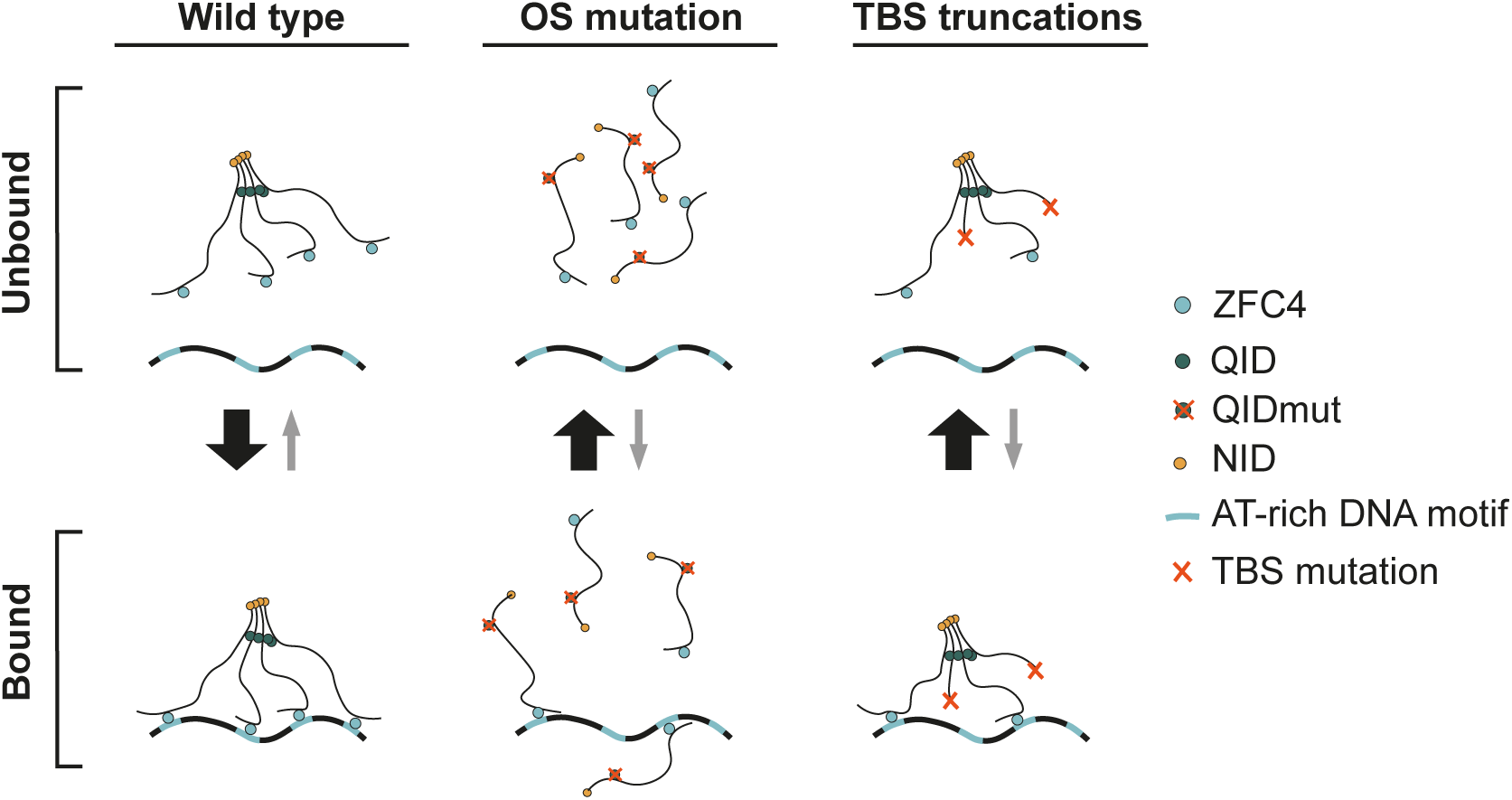
SALL protein multimerisation is essential for efficient DNA binding and relevant for the molecular disease mechanism of Okihiro and Townes-Brocks syndromes. Schematics of the importance of SALL protein multimerisation for its physiological function and its relevance in the disease mechanisms of OS and TBS respectively. The QID is shown as a green dot (crossed out for QIDmut) and the NuRD interaction domain (NID) is coloured in yellow. The DNA-binding domain ZFC4 is shown as a blue dot and AT-rich motifs in the DNA as blue stretches. TBS-causing frameshift and nonsense mutations leading to premature stop codons in SALL1 are shown as red crosses.

## Discussion

In this study, we defined a Q-rich Interaction Domain (QID) in SALL proteins that is responsible for tetramerisation and high DNA-binding avidity, and explored both its physiological and pathological relevance. We demonstrate that tetramerisation of a relatively weak SALL4 DNA-binding domain^9,11^ greatly increases DNA-binding avidity, leading to efficient chromatin binding *in vivo*. In support of this hypothesis, disruption of the SALL4 multimerisation domain (QIDΔ and QIDmut) abolished almost all binding to the genome and resulted in a strongly deregulated transcriptome, similar to a full SALL4 knockout. We observed a reduction in SALL4 protein stability in these cell lines (≈50% reduction by western blot compared to WT cells), but this cannot explain such a dramatic loss of SALL4 function, as a HET S4KO control line expressing SALL4 to a similar level did not recapitulate chromatin binding defects. Additionally, proteomic analyses revealed that monomeric SALL4 (QIDΔ and QIDmut) interacts less strongly with components of the NuRD co-repressor complex by ∼4-fold compared to WT SALL4. This suggests that SALL4 tetramerisation also increases avidity for co-repressor subunits via protein-protein interactions. It may not be coincidental that each NuRD complex is thought to contain four molecules of RBBP4^49,50^, the target protein for the SALL4 NuRD interaction domain. Taken together, our data identify the QID as a critical functional domain of SALL4.

SALL4 was detected in a screen for proteins that recognise multiple A/T-motifs^8^ and might therefore be a reader of DNA base composition^51^. Subsequent experiments have strengthened this conclusion by showing that SALL4 lacking the ZFC4 DNA binding domain causes preferential up-regulation of ESC genes in proportion to their A/T-richness^8^. ESCs with this mutation (like *Sall4* knockout ESCs) differentiate promiscuously and upregulate genes that are activated naturally during the differentiation process, suggesting that the presence of wildtype SALL4 stabilises the stem cell state^8,14^. While the targeting of the SALL4 DNA binding motif to runs of As and Ts could explain its preference for A/T-rich DNA, it is apparent that tetramerisation potentially enhances affinity. If we hypothesise that each ZFC4 in the tetramer needs to contact an A/T motif for the interaction to become stable, then there will be a requirement for 4 accessible A/T motifs within the genomic span reachable by one tetramer. Leaving aside the complicating effects of nucleosomes on binding, which are currently unknown, it is evident that fluctuations in base composition will affect A/T motif clustering, as the average frequency of motifs will be acutely sensitive to base composition.

The minimal SALL4 multimerisation domain mapped here is longer than the 22 amino acids Q-rich region previously proposed to form the interaction domain^31^. These additional residues may be required for the stability of the helix, but also for the specificity of the interaction as there is one more charged residue in the N-terminus of the domain (E211), which could potentially mediate salt-bridge formation^52^. In addition to this, the proline in position 3 could serve as a helix-initiator and cap it at the N-terminus, as has been described for other alpha-helices^53^. While the SALL4 QID does not follow the classical spacing of polar and non-polar residues normally found in a coiled coil^54^, it has some sequence and predicted structural similarities to a Q-rich region found in HDAC4 that is also known to promote homotetramer formation^55^. These shared features hint at the possibility that Q-rich regions might belong to a set of protein-protein interaction domains more diverse than the much studied classical leucine-zippers which follow the defined spacing^56^. Conservation of the QID in all SALL family proteins in both vertebrate and invertebrate species along with our IP-MS data predicts that these 33 amino acids are also necessary and sufficient for multimerisation in SALL1, 2 and 3.

While the QID might mediate multimerisation of SALL1, 2, 3 and 4, the frequency and composition of the entire population of SALL tetrameric complexes remains unknown. We anticipate that the level of expression of the different members of the SALL protein family in a given cellular context is a critical determinant. In mouse ESCs, SALL4 is predominant, and therefore a proportion of SALL protein complexes in this cell type will probably correspond to SALL4 homotetramers. In contrast, differentiated cell types such as kidney cells^57^, microglia^58^ or neurons^59^ express high levels of SALL1, 2 and 3, which is likely to result in increased heterotetramer formation and could be functionally relevant for the regulation of developmental processes. Future work will examine the relative expression levels of SALL proteins in embryonic and adult tissues as well as the functional heterogeneity associated with these assemblies.

Heteromultimerisation of truncated SALL1 with other SALL proteins has been proposed as responsible for the dominant-negative effect of TBS-causing mutations^29^. However to date, the molecular explanation underlying phenotypic evidence in animal models^25,29^ and phenotype-genotype correlations in TBS patients^47,60,32^ remained unclear. We now suggest that “poisoning” of tetrameric SALL protein complexes by TBS-causing SALL1 truncations lacking DNA-binding domains but retaining the QID could be responsible for pathogenesis. While this mechanism would explain the majority of pathogenic mutations localising within a hotspot, around 13% of TBS-causing mutations would cause SALL1 truncation before or within the QID. However, phenotypic data is only available for two of these cases, one patient with the central symptomatic triad of TBS (313del^47^) and the other (420del^28^) only presenting milder symptoms including dysplastic ears, misplaced finger-like thumbs and reduced kidney clearance. Thus, it cannot be excluded that many of those patients are diagnosed with TBS despite having milder symptoms compared to typical cases bearing mutations that truncate SALL1 after the QID.

Our findings may also have clinical relevance in cancer. SALL4 is overexpressed in many cancers with poor prognosis^61–67^ and there is some evidence that it may act as an oncogene^68^. Accordingly, several strategies have been investigated to attenuate SALL4 function in cancer cells, including targeting of SALL4 NID^5^ or C2H2 ZFCs^12^. However, these approaches risk causing off-target effects as other TFs contain highly similar domains. In contrast, the SALL QID could represent a more specific target that warrants experimental investigation.

While increased avidity through multimerisation is well known (e.g. haemoglobin^69,70^), it remains less well characterised in the context of TF biology^71^. The presence of *cis*-regulatory modules in invertebrate and vertebrate genomes that contain homotypic clusters of transcription factor binding sites^72^ raises the possibility that the importance of multivalent binding by TFs may be underappreciated. Few studies have conclusively shown that TF multimerisation is a pre-requisite for efficient DNA-binding (e.g. SOX6^73^), but more recent works on BTB domain proteins^74^ and FOXP3^75^ have suggested that higher-order assemblies of these TFs are needed for their cellular functions. Together, these findings raise the possibility that defined, higher-order multimerisation - often mediated by Q-rich regions^76^ - is a more general strategy used by TFs to fine-tune DNA binding and regulatory activity, but one that remains underexplored. Our study therefore establishes a framework for considering multimerisation as a central, and perhaps widespread, principle in transcription factor biology.

## Materials and Methods

### EXPERIMENTAL MODEL AND SUBJECT DETAILS

#### Cell culture

All cell lines used in this study correspond to embryonic stem cells (ESCs) from *Mus musculus*. The WT line used in this study is JU09, which was derived from 129/Ola E14Tg2a cells^77^. Furthermore, a *Sall4* knockout line (S4KO) was kindly provided by Dr Brian Hendrich^14^ and an E14/T WT line for overexpression of transfected pPYCAG constructs was obtained from Prof. Ian Chambers^78,79^. Other cell lines (QIDΔ, QIDmut, new S4KO, HET S4KO, S4-mScarlet3) were derived in this study by genetic engineering of JU09 ESCs, detailed in the section below.

All ESC lines were cultured at 37°C and 5% CO_2_ on gelatine-coated dishes. For general maintenance, cells were grown in Glasgow minimum essential medium (GMEM; GIBCO ref. 11710035) supplemented with 15% fetal bovine serum (FBS; GIBCO ref. 10270106, batch tested), 100U/mL leukemia inhibitory factor (LIF; Sigma Aldrich ref. ESG1107, batch tested), 1x L-glutamine (GIBCO ref. 25030024), 1x MEM non-essential amino acids (GIBCO ref. 11140035), 1 mM sodium pyruvate (GIBCO ref. 11360070) and 0.1 mM beta-mercaptoethanol (GIBCO ref. 31350010).

To transiently transfect ESCs for overexpression or genetic manipulation, Lipofectamine 3000 (Invitrogen ref. L3000015) was used following manufacturer’s instructions. For selection of transfected cells, the culture medium was supplemented with puromycin (GIBCO ref. A1113803) to a final concentration of 10 µg/mL.

#### Animal work

The Sall4 L221P (QIDmut) mouse line was generated by injecting CRISPR/Cas9-targeted heterozygous ESCs (see section below) into mouse blastocysts. Male and female chimeras were bred with C57BL/6J wild-type mice from seven weeks old and germline transmission of the targeted allele was identified by coat colour and confirmed by PCR (see Methods S1; Genotyping primers) and Sanger sequencing. Heterozygous offspring were interbred to obtain homozygotes and genotyping was routinely performed on all offspring by PCR with BbVCI-based restriction fragment length polymorphism (RFLP) analysis, which specifically targeted the L221P allele. Heterozygotes showed no observable phenotype in either sex at any age, but they were born at sub-Mendelian ratios. Statistical analysis was performed in GraphPad/Prism (v9) using a Fisher’s Exact test contrasting the born genotypes to the expected ones if inheritance of the allele was Mendelian. All mice were bred and maintained in SPF conditions at the University of Edinburgh. Procedures were conducted by UK Home Office–licensed staff in accordance with the Animal and Scientific Procedures Act 1986 and approved by the local Animal Welfare and Ethical Review Board. Mice were kept on a 12-hour light/dark cycle with ad libitum food and water, housed in individually ventilated cages with wood chip bedding, tissue nesting and environmental enrichment, in groups of up to five animals. Mutant and wild-type littermates were housed in the same cages.

### METHOD DETAILS

#### Genetic manipulations of ESCs

To introduce mutations or tags into the endogenous *Sall4* locus (QIDΔ, QIDmut, S4KO, HET S4KO, S4-mScarlet3), WT JU09 ESCs were edited using CRISPR/Cas9^80^. Guide RNAs were designed near the target site (https://benchling.com) and cloned into Cas9/gRNA co-expression plasmids (pX458 containing EGFP and pX459 V2.0 containing puromycin resistance cassette, AddGene refs. 48138 and 62988). Single-stranded repair DNA templates (ssDNAs) for homologous recombination were obtained from Integrated DNA Technologies and contained mutations of interest as well as PAM mutations to prevent further cutting of the repaired allele. The targeting vector used for the generation of the S4-mScarlet3 cell line was cloned by Gibson assembly using the NEB HiFi assembly kit (NEB ref. E2621L) and following manufacturer’s instructions (see Methods S1; CRISPR oligos & repair templates). ESCs (1x10^6^ cells) were transfected with one Cas9/gRNA plasmid for point mutations and tagging or two for deletions, along with 10 nmol ssDNA template or 0.3 pmol targeting vector. Cells were selected with 10 µg/mL puromycin and/or sorted for GFP/mScarlet3 fluorescence by flow cytometry depending on the selection cassette contained in transfected plasmids. Single cells were sorted by Fluorescence-Activated Cell Sorting (FACS) into 96-well plates to obtain ESC clones, which were expanded and from which genomic DNA was extracted. Genotyping of desired mutations was confirmed by PCR (see Methods S1; Genotyping primers) and Sanger sequencing.

To generate doxycycline-inducible overexpression (OE) cell lines (SALL4 WT OE, QIDΔ OE or QIDmut OE), *Sall4* cDNA constructs were randomly integrated into S4KO ESCs using an adapted PiggyBac (PB) transposon system (see previous publication for details^8^). 1x10^6^ S4KO ESCs were transfected with two PiggyBac vectors and a third plasmid expressing hyperactive PB transposase^43^. One of the piggyBac vectors was carrying *Sall4* under a minimal Dox-inducible promoter and contained a hygromycin B selection cassette, while the other carried the TetON transactivator under a constitutive promoter as well as a zeocin selection cassette. After 48 hours, transfected cells were selected with 200 µg/mL hygromycin B and 100 µg/mL zeocin for 10 days. Surviving cells were pooled and frozen in aliquots prior to any doxycycline induction. To induce SALL4 overexpression, cells were treated for 48 hours with freshly prepared 1 µg/mL doxycycline.

#### Immunoprecipitation

To prepare nuclear protein extracts for immunoprecipitation and subsequent analysis of co-immunoprecipitated proteins by western blot, ESCs (5-10 million) were washed in PBS, detached using TrypLE (GIBCO ref. 12604013) and collected by centrifugation (300 x *g* for 5 minutes) in 15 mL tubes. The supernatant was removed and pellets were resuspended in 1 mL lysis buffer (20 mM HEPES pH 7.5, 10 mM NaCl, 1 mM MgCl2, 0.1% (v/v) Triton X-100, freshly supplemented with 1x protease inhibitor cocktail (PIC, Roche ref. 11873580001) and 0.5 mM DTT. Following 20 minutes of incubation on ice, nuclei were collected by centrifugation (200 x *g* for 10 minutes at 4°C). The supernatant containing cytoplasmic proteins was discarded, and nuclei were resuspended in 1 mL of fresh lysis buffer freshly supplemented with 250 U of benzonase nuclease (Millipore ref. E1014). Following five minutes incubation at room temperature, 5 M NaCl was added to samples to achieve a final concentration of 150 mM. Samples were then incubated on a rotating wheel at 4°C for 30 minutes, followed by a 30-minute centrifugation at 15,700 x *g* and 4°C. The resulting supernatants containing solubilised nuclear proteins were collected into fresh tubes and input material was prepared by boiling 40 μL of nuclear protein extracts at 90°C for five minutes in 40 μL Laemmli buffer (Sigma Aldrich ref. S3401). Nuclear extracts were immediately used for immunoprecipitation or stored at -70°C.

For immunoprecipitation of Flag-tagged constructs, anti-FLAG M2 magnetic beads (Millipore ref. M8823) were used. Beads were washed three times in 1 mL PBS, added to nuclear protein extracts (30 µL per IP) and incubated for two hours at room temperature on a rotating wheel. Beads were then washed three times in 1 mL PBS and finally boiled in 30 µL Laemmli buffer for five minutes at 90°C to elute the immunoprecipitated material.

Endogenous SALL4 was immunoprecipitated with 5 µg of rabbit anti-SALL4 polyclonal antibody (Abcam ref. ab29112) overnight on a rotating wheel at 4°C. The next day Protein A Sepharose beads (Cytiva ref. 17528001), which were pre-blocked with 0.5 mg/mL BSA and washed three times in 1 mL lysis buffer, were added to the samples (50 µL per IP) and incubated for two hours on a rotating wheel at 4°C. Beads were subsequently washed five times in 1 mL lysis buffer with centrifugation steps (one minute at 400 x *g* and 4°C) to collect beads between each wash. Finally, beads were resuspended in 30 µL Laemmli buffer and boiled for five minutes at 90°C to elute immunoprecipitated proteins off the beads.

#### Western blot

Protein samples were loaded into precast polyacrylamide gels (4-15% or 4-20%, Bio-Rad refs. 4561095, 4561083 and similar) together with a molecular weight marker (Li-Cor ref. 928-60000). Gels were run in SDS running buffer for one hour at 200 V to size-separate the proteins. Proteins were then transferred onto a nitrocellulose membrane at 100 V for one hour. The membrane was then blocked in 10% milk (w/v) dissolved in PBS-Tween (1x PBS, 0.1% v/v Tween 20) for one hour at room temperature. Primary antibodies (see Methods S1; Antibodies) were diluted into 5% milk (w/v) dissolved in PBS-Tween and incubated with the membrane for one hour shaking at room temperature or overnight at 4°C. Membranes were submitted to four five-minute washes in PBS-Tween, and incubated with fluorescently-labelled secondary antibodies (see Methods S1; Antibodies) diluted in 5% milk dissolved in PBS-Tween for one hour at room temperature under constant shaking. Following four five-minute washes in PBS-Tween, fluorescent signals were detected and imaged on the Odyssey CLx instrument. Intensities within specific protein bands were quantified using the Image Studio software (v5.2.5). To quantify protein signal, fluorescence was normalised to a control corresponding to the immunoprecipitated protein in the case of co-IP, or Histone H3 loading control in the case of inputs. Data was scaled to a positive control for visualisation. Calculations and visualisations were done using Microsoft Excel (v16.80) and GraphPad/Prism (v9). Each co-IP experiment was repeated for at least three times to obtain technical replicates.

#### Mass Spectrometry

##### Immunoprecipitation

2.4 million cells were harvested by centrifugation into Protein LoBind tubes, washed with PBS, resuspended in 1 mL ice-cold 1x NE buffer (20 mM HEPES, 10 mM NaCl, 1 mM MgCl₂, 0.1% (v/v) Triton X-100, 1x PIC) and incubated on ice for 20 minutes. Nuclei were pelleted by centrifugation (500 x *g*, 10 minutes, 4°C) resuspended in 1 mL of fresh NE buffer and treated with 125 U of benzonase nuclease (Millipore ref. E1014) for five minutes at room temperature. 5 M NaCl was then added to a final concentration of 150 mM in each sample, followed by an incubation on a rotating wheel for 30 minutes at 4°C. Samples were centrifuged (16,000 x *g*, 15 minutes, 4°C) to remove debris and supernatants were transferred into fresh tubes. 500 µL Protein G Dynabeads (Invitrogen ref. 10003D) were washed once in lysis buffer (50 mM Tris-HCl pH 8, 150 mM KCl, 0.1% (v/v) Triton X-100) and incubated with 50 µg SALL4 antibody (Santa Cruz ref. sc-101147) in 2 mL lysis buffer for one hour at room temperature on a rotating wheel. Beads were then washed three times in lysis buffer (10 minutes incubation between each wash), resuspended in 10 mL borate buffer (40 mM boric acid, 40 mM sodium tetraborate decahydrate) containing 20 mM DMP (Thermo Scientific ref. 21666) for crosslinking and incubated for 30 minutes at room temperature on a rotating wheel. After a five-minute wash in lysis buffer, beads were resuspended in 1 mL lysis buffer. For IP, 100 µL crosslinked beads (equivalent to 5 µg antibody) were added to 1 mL nuclear extract and incubated for two hours at 4°C on a rotating wheel. Beads were then washed twice with 1 mL lysis buffer and once with 100 µL lysis buffer. Proteins were finally eluted from the beads by incubation in 50 µL of 0.1% (w/v) Rapigest (VWR ref. WATE186001861) dissolved in 50 mM Tris-HCl pH 8 for 15 minutes at 60°C. Eluates were transferred into fresh tubes and used for filter-aided sample preparation (FASP).

##### Filter-aided sample preparation (FASP)

5 µL of freshly prepared 1 M DTT (dissolved in 100 mM Tris-HCl pH 8) was added to each sample, followed by incubation on a thermomixer (80°C, 500 rpm, 15 minutes). Samples were cooled down at room temperature, mixed with 100 µL UBB (8 M urea, 100 mM Tris-HCl pH 8), loaded onto Vivacon 500 spin columns (Sartorius ref VN01H21) and centrifuged at 10,000 x *g* for 20 minutes. Columns were then treated with 100 µL UBB + IAA (8 M urea, 100 mM Tris-HCl pH 8, 100 mM iodoacetamide), briefly shaken (600 rpm, one minute) and incubated for 30 minutes at room temperature in the dark. After centrifugation (13,500 x *g*, 15 minutes), columns were washed with 100 µL UBB and twice with 75 µL ABC (50 mM ammonium bicarbonate in ddH₂O) with centrifugation steps between each wash to ensure the membranes were dry. Columns were transferred into fresh collection tubes and 100 µL trypsin solution (0.6 µg trypsin (Thermo Scientific ref. 90057, dissolved in 100 µL ABC buffer) was added to each sample. Columns were shaken (600 rpm, one minute), sealed with parafilm and incubated overnight at 37°C. The following day, peptides were eluted by centrifugation (10,000 x *g*, 15 minutes), followed by an additional elution with 75 µL ABC. Combined eluates (100–150 µL) were acidified to pH ∼1–2 with 5 µL 10% TFA (Sigma Aldrich ref. T6508-100ML). Samples were loaded onto stage tips^81^ and centrifuged at 700 x *g* for 10–20 minutes until all liquid passed through. Lastly, samples were washed with 75 µL 0.1% TFA and centrifuged at 1,000 x *g* for six minutes before being stored at –20°C until mass spectrometry analysis.

##### LC-MS/MS analysis

LC-MS analyses for the IPs were performed on an Orbitrap Fusion™ Lumos™ Tribrid™ Mass Spectrometer (Thermo Fisher Scientific, UK) and for the nuclear extracts on Orbitrap Exploris 480™, both coupled on-line, to an Ultimate 3000 HPLC (Dionex, Thermo Fisher Scientific, UK). Peptides were separated on a 50 cm (2 µm particle size) EASY-Spray column (Thermo Scientific, UK), which was assembled on an EASY-Spray source (Thermo Scientific, UK) and operated constantly at 50°C. Mobile phase A consisted of 0.1% formic acid in LC-MS grade water and mobile phase B consisted of 80% acetonitrile and 0.1% formic acid. Peptides were loaded onto the column at a flow rate of 0.3 μL/min and eluted at a flow rate of 0.25 μL/min according to the following gradient: 2 to 40% mobile phase B in 150 min and then to 95% in 11 min. Mobile phase B was retained at 95% for 5 min and returned back to 2% a minute after until the end of the run (190 min).

For the IPs, survey scans were recorded at 120,000 resolution (scan range 350-1500 m/z) with an ion target of 4.0e5, and injection time of 50ms. MS2 was performed in the ion trap at a rapid scan mode, with ion target of 2.0e4 and HCD fragmentation^82^ with normalized collision energy of 27. The isolation window in the quadrupole was 1.4 Thomson. Only ions with charge between 2 and 7 were selected for MS2. Dynamic exclusion was set at 60 s. The MaxQuant software platform^83^ (version 1.6.1.0) was used to process the raw files and search was conducted against the Mus musculus protein database (released in July 2017), using the Andromeda search engine^84^. For the first search, peptide tolerance was set to 20 ppm while for the main search peptide tolerance was set to 4.5 pm. Isotope mass tolerance was 2 ppm and maximum charge to 7. Digestion mode was set to specific with trypsin allowing maximum of two missed cleavages. Carbamidomethylation of cysteine was set as fixed modification and oxidation of methionine was set as variable modifications. Additionally, methylation of lysine and arginine, di- and tri-methylation of lysine were also set as variable modifications. Label-free quantitation analysis was performed by employing the MaxLFQ algorithm as previously described^85^. Absolute protein quantification was performed as previously described^86^. Peptide and protein identifications were filtered to 1% FDR.

For the nuclear protein extracts, MS1 scans were recorded at 120,000 resolution (scan range 350-1650 m/z) with an ion target of 5.0e6, and injection time of 20 ms. MS2 Data Independent Acquisition (DIA) was performed in the orbitrap at 30,000 resolution with a scan range of 350-1200 m/z, maximum injection time of 55 ms and AGC target of 3.0e6 ions. We used HCD fragmentation^82^ with stepped collision energy of 25.5, 27 and 30. We used variable isolation windows throughout the scan range ranging from 10.5 to 50.5 m/z. Shorter isolation windows (10.5-18.5 m/z) were applied from 400-800 m/z and then gradually increased to 50.5 m/z until the end of the scan range. The default charge state was set to 3. Data for both survey and MS/MS scans were acquired in profile mode. The DIA-NN software platform^87^ (version 1.8.1). was used to process the raw files and search was conducted against the Mus musculus complete/reference proteome (Uniprot, released in July 2017). Precursor ion generation was based on the chosen protein database (automatically generated spectral library) with deep-learning based spectra, retention time and IMs prediction. Digestion mode was set to specific with trypsin allowing maximum of two missed cleavages. Carbamidomethylation of cysteine was set as fixed modification. Oxidation of methionine, and acetylation of the N-terminus were set as variable modifications. The parameters for peptide length range, precursor charge range, precursor m/z range and fragment ion m/z range as well as other software parameters were used with their default values. The precursor FDR was set to 1%. Statistical analysis was performed by Perseus software^88^ (version 1.6.2.1). Data was plotted using Microsoft Excel (v16.80) and GraphPad/Prism (v9).

#### RNA-seq

##### Sample preparation for RNA-seq of transiently transfected ESCs

200,000 pre-seeded WT ESCs (E14^77^) were transfected with equimolar amounts of GFP-tagged expression constructs (GFP, S1N, S1Nmut) using Lipofectamine 3000 and following manufacturer’s instructions. The day after transfection, cells were allowed to recover for seven hours in normal ESC culture medium (see composition above), before supplementation with 10 µg/mL puromycin for 48 hours (GIBCO ref. A1113803). Cells were assessed for GFP expression by microscopy at the end of the selection period and allowed to recover in ESC media without puromycin for another 20 hours before being harvested. Cells were counted to collect exactly 600,000 cells by centrifugation (300 x *g* for five minutes) and those cell pellets were then directly lysed in lysis buffer (Qiagen ref. 74134) for subsequent RNA extraction following manufacturer’s instructions (including QIAshredder (Qiagen ref. 79656) and gDNA eliminator columns). Purified RNA was quantified using the Qubit RNA High Sensitivity Assay kit (Invitrogen ref. Q32852) and stored at -70°C until library preparation. In addition, 150,000 cells were harvested from the same samples and lysed in 45 µL Laemmli buffer for quality control to check SALL1/4 protein levels by western blot. The experiment was repeated three times to obtain triplicates and untransfected WT cells as well as other control ESCs (S4KO HET and S4KO - for RT-qPCR only) were grown and harvested simultaneously to transfected samples, but were not subjected to the selection process.

##### RNA extraction from genetically modified cell lines

ESCs were grown for 96 hours in N2B27 + 2i/LIF medium to ensure a pluripotent phenotype for all cell lines, followed by 96 hours in standard culture condition (Serum/LIF) prior to harvesting. 2.5 million ESCs of each genotype were counted and resuspended in 500 µL TRIzol reagent (Invitrogen ref. 15596026). RNA was subsequently extracted using a Direct-zol RNA Miniprep Plus kit (Zymo Research ref. R2070) following manufacturer’s instructions including the DNAseI treatment. RNA was eluted in 60 µL DNase/RNase-free water, quantified using the Qubit RNA High Sensitivity Assay kit (Invitrogen ref. Q32852) and stored at -70°C. This experiment was repeated four times to obtain quadruplicates.

##### Library preparation

1 µg of total RNA was used as input material for each sample. Libraries were prepared using the KAPA RNA HyperPrep Kit with RiboErase HMR (v2.17, Roche ref. 08098131702) and the manufacturer’s protocol was followed with the following adjustments: In step 6 (RNA elution, Fragmentation and Priming) the sample mix was incubated at 94°C for seven minutes to obtain ≈200 bp fragments. Furthermore, the KAPA dual-indexed adapters (Roche ref. 08278555702) used in step 9 (Adapter Ligation) were diluted to 7 µM with the provided dilution buffer. Lastly, the adapter-ligated DNA was amplified for a total of seven PCR cycles in step 12 (Library amplification). The final libraries were quantified on the Qubit system (Invitrogen ref. Q32851) and library size was analysed using the 2100 bioanalyzer instrument and High-Sensitivity DNA Analysis kit (Agilent ref. 5067-4626). Barcoded libraries passing these quality-control steps were pooled in equimolar ratios and sequenced using the Illumina NovaSeq platform (paired-end, Novogene or Genewiz UK).

#### RT-qPCR

1 µg of purified RNA was used as input material for each sample and reverse-transcribed with SuperScript IV and random hexamer primers (Invitrogen ref. 18091050) following manufacturer’s instruction, including the generation of a negative control without reverse transcriptase (no RT) for each replicate experiment in order to detect gDNA contamination. The obtained cDNA libraries were diluted 1:20 in water and 4.5 µL of material was used for qPCR reaction using the Takyon SYBR Mastermix (Eurogentech ref. UF-NSMT-B0701) with appropriate primer pairs (see Methods S1; RT-qPCR primers), following manufacturer’s instructions (Roche Lightcycler 480 instrument, Takyon FAST cycling parameters). For each sample, PCR reactions were performed in technical triplicate wells of a 384-well plate, and average signal was used for analysis. For each primer pair melting curves were generated and analysed to verify the production of single DNA species. The data was further analysed using Microsoft Excel (v16.80) and visualised with GraphPad/Prism (v9), where all transcript levels were normalised to TATA-binding protein (TBP) and expressed relative to WT.

#### Spontaneous differentiation in N2B27 and alkaline phosphatase staining

This experiment followed previously published protocols^89,90^. To ensure a pluripotent phenotype before the assay and for colony growth after it, cells were grown in N2B27 medium containing a 1:1 mix of Advanced DMEM/F-12 (GIBCO ref. 12634010) and Neurobasal (GIBCO ref. 21103049) supplemented with 1x L-Glutamine (GIBCO ref. 25030024), 1x MEM non-essential amino acids (GIBCO ref.11140035), 0.5x N-2 supplement (GIBCO ref. 17502048), 0.5x B-27 supplement (GIBCO ref. 17504044) and 0.1 mM beta-mercaptoethanol (GIBCO ref. 31350010), as well as ‘‘2i’’ inhibitors^91^ (1mM PD0325901 (Axon ref. 1408) and 3mM CHIR99021 (Axon ref. 1386)) and 100U/mL LIF were added.

To spontaneously differentiate ESCs, cell lines were cultured in N2B27 medium (see composition above) without the addition of 2i inhibitors and LIF.

S4KO ESCs with *Sall4* transgenes were grown in N2B27 + 2i/LIF for 4 days, washed with PBS (GIBCO ref. 14190094), detached using Accutase (GIBCO ref. A1110501) and 100,000 cells were plated on gelatine-coated six well plates in serum-free N2B27 medium supplemented with 1 µg/mL freshly prepared doxycycline (Sigma Aldrich ref. D9891) to induce transgene expression. Cells were allowed to spontaneously differentiate for four days with a medium change on day two. After differentiation, cells were collected and seeded at clonal density (4,000 cells per well) in technical triplicates on Matrigel-coated (diluted 1:100 in cold DMEM; Corning ref. 354234, GIBCO ref. 41966-029, pre-coated for one hour) six well plates containing N2B27 medium + 2i/LIF to recover ESC colony growth. After 7-10 days, colonies were fixed and stained for alkaline phosphatase (AP) expression using a commercial kit and following manufacturer’s instructions (Sigma Aldrich ref. 86R-1KT). Plates were scanned and AP+ colonies were quantified using a custom-made ImageJ plugin (see quantification and statistical analysis; Automated quantification of AP-staining section). As a control, undifferentiated cells were plated at clonal density (600 cells/well) and stained for AP after 10 days of culture in N2B27 +2i/LIF.

#### Immunofluorescence

150,000 cells were seeded on gelatine-coated microscopy dishes (Ibidi ref. 80286). At 50-80% confluency, cells were washed with PBS, fixed with 1 mL of 4% PFA for 10 minutes at room temperature, washed with PBS and permeabilised with 1 mL permeabilisation buffer (1x PBS, 0.3% (v/v) Triton X-100) for 10 minutes at room temperature. Afterwards, cells were blocked with 0.5 mL blocking buffer (1x PBS, 0.1% (v/v) Triton X-100, 1% (w/v) BSA, 3% (v/v) goat serum) for one hour shaking at room temperature. Samples were incubated with primary antibodies diluted in blocking buffer (see Methods S1; Antibodies) for one hour at room temperature or overnight at 4°C, followed by four five minutes washes with 1 mL PBS-Triton (1x PBS + 0.1% (v/v) Triton X-100). Samples were subsequently incubated with fluorescently-labelled secondary antibodies (see Methods S1; Antibodies) diluted in blocking buffer for one hour at room temperature, with slides wrapped in aluminium foil due to light sensitivity. After four five-minutes washes with 1 mL PBS-Triton, DNA was stained with 1 µg/mL DAPI diluted in PBS (Sigma Aldrich ref. MBD0015) for five minutes shaking at room temperature. Finally, samples were washed with 1 mL PBS and either stored in PBS (short-term) or mounted with 2-3 drops of ProLong glass antifade mountant (Invitrogen ref. P36980) and coverslips (long-term) at 4°C. Immunofluorescently labelled cells were imaged on the Zeiss LSM 880 Airyscan confocal microscope using z-stacks to capture entire cells. Images were processed using the ZEN software and analysed using ImageJ^92^. For each replicate experiment, cells were individually assessed for SALL4 co-localisation with DNA foci and the percentage of cells with foci was calculated. Each replicate experiment included WT and S4KO controls. Calculations were done using Microsoft Excel (v16.80) and visualisation was performed using GraphPad/Prism (v9).

#### Live-cell imaging

150,000 SALL4-mScarlet3 ESCs were seeded on gelatine-coated ibidi dishes (80286) and imaged one day after transfection with SALL1-GFP expression constructs using Lipofectamine 3000 (Invitrogen ref. L3000015) and following manufacturer’s instructions. Medium was changed 5 min prior to imaging and two drops of the NucBlue/Hoechst reagent (Invitrogen ref. R37605) was added to each well to stain DNA in live cells. Samples were imaged using the Zeiss LSM 880 Airyscan confocal microscope with 37°C and 5% CO_2_ incubation. Z-stacks were acquired and images were processed using the ZEN software and further analysed using a custom-made ImageJ plugin (see quantification and statistical analysis; Quantification of foci/nucleoplasm ratios in live-cell imaging section).

### CUT&RUN

#### Extraction of target DNA

CUT&RUN was performed using the EpiCypher CUTANA ChIC/CUT&RUN kit (v3.1, Stratech ref. 14-1048-EPC) following manufacturer’s instructions. In short, 500,000 cells per genotype were used and bound to pre-activated ConA beads, followed by washes and permeabilisation in a buffer containing 0.01% (v/v) digitonin as well as 2 mM EDTA. Permeabilised cells were then incubated overnight in buffer containing 0.5 µg of antibodies of interest (see Methods S1; Antibodies). The following day, cells were washed to remove unbound antibodies, then incubated with 1x pAG-MNase. The digestion was activated by addition of CaCl₂ (final concentration 2 mM) and incubation for two hours at 4°C. The reaction was stopped and 0.5 ng *E. coli* DNA spike-in was added to each sample for normalisation. Finally, released DNA fragments were purified using the provided spin columns, quantified on a Qubit system (Invitrogen ref. Q32851) and stored at -20°C until library preparation.

#### Library preparation

Libraries for next-generation sequencing were generated using the KAPA HyperPrep Kit (v7.19, Roche ref. 07962347001) with KAPA dual-indexed adapters (Roche ref. 08278555702) following the manufacturer’s protocol with some modifications to account for the low yield and small size of DNA fragments generated by CUT&RUN. For each reaction, 5 ng of DNA was used as input. End-repair and A-tailing were carried out at 20°C for 30 minutes and 58°C for 45 minutes (instead of 65°C) to prevent melting of small fragments. Adapters were diluted at 1.5 µM and ligated with DNA fragments for 30 minutes (instead of 15 minutes) at 20°C to improve conversion efficiency. DNA was subjected to an additional selection with 1.1x KAPA beads during cleanup post-ligation. Library amplification was performed with cycling conditions at the maximum of the advised range to enrich for small fragments. Amplified libraries were subjected to an additional selection with 1.1x KAPA beads during final cleanup and eluted in 22 µL elution buffer (EB, 10 mM Tris-HCl pH 8.5 in water). Libraries were quantified using the Qubit system (Invitrogen ref. Q32851) and library sizes were assessed with an Agilent 2100 Bioanalyzer and High-Sensitivity DNA kit (Agilent ref. 5067-4626). Libraries passing quality control were pooled in equimolar ratios and sequenced on Illumina NovaSeq platforms by Novogene or Genewiz UK (paired-end sequencing).

#### Recombinant protein purification

Bacterial expression plasmids (His-GFP and His-MBP) were gifted by Dr Marcus Wilson’s laboratory (Edinburgh UK) and used for sub-cloning SALL4 constructs by Gibson assembly using the HiFi assembly kit (NEB ref. E2621L). Plasmids were transformed into BL21 Star competent cells (Invitrogen ref. C601003) and a starter culture was grown in liquid LB supplemented with the appropriate antibiotic selection overnight shaking at 37°C. 1 mL of the starter culture was then diluted into 100 mL of Magic Medium (Invitrogen ref. K6803) supplemented with antibiotics and grown shaking at 37°C until exponential phase (OD_600_ = 0.6). 0.4 mM IPTG (Fisher Scientific ref. V3955) was added to the medium to initiate protein production, and cells were incubated overnight shaking at 18°C. The next day, the bacteria were harvested by centrifugation (2,095 x *g*, 15 minutes, 4°C) and pellets were either frozen at -70°C or directly used for protein extraction and purification.

To purify the minimal SALL4 QID peptide, cell pellets were resuspended in cold, filtered lysis buffer (250 mM NaCl, 25 mM Tris pH 7.5; 10 mL per gram of wet cell pellet) supplemented with 1x protease inhibitor cocktail (PIC, 1/100 ml). Cells were lysed using a Constant Systems cell disruptor (benchtop TS 1.1 kW, 25 kpsi, 6°C) and the sample was centrifuged (39,000 x *g*, 30 minutes, 4°C) to pellet debris. The supernatant containing soluble proteins was filtered and supplemented with 20 mM imidazole (VWR ref. 286874D) to reduce nonspecific binding of other bacterial proteins. His-tagged SALL4 constructs were purified on an ÄKTA pure chromatography system using an IMAC column (Cytiva ref. 17092102), washed with 20 column volumes (CV) of 20 mM imidazole and 10 CV of 30 mM imidazole, then eluted in a block with 500 mM imidazole. Fractions were collected and analysed by SDS-PAGE, Coomassie staining and immunoblotting. Fractions containing protein of the correct size were pooled and desalted over a HiPrep 26/10 column (Cytiva ref. 17508701) to remove imidazole. Proteins were digested overnight with TEV protease (1 mg TEV/100 mg protein,) and cleavage of the affinity tag was assessed by SDS-PAGE. The sample was then passed over an IMAC column again, which bound His-GFP, and the flow-through containing untagged SALL4 QID was collected. SALL4-containing fractions were pooled and concentrated in a spin concentrator with the appropriate molecular weight cut-off (10,000 x *g*, 10 minutes, 4°C). Finally, the sample was purified by gel filtration (Superdex 75 pg, HiLoad 16/600, Cytiva ref. 28989333), and fractions containing SALL4 were identified by SDS-PAGE, pooled, concentrated again and frozen in aliquots at –70°C.

To purify SALL4 QID-ZFC4 fusion peptides, cell pellets were resuspended in cold, filtered lysis buffer (500 mM NaCl, 50 mM Tris pH 7.5, 10% (v/v) glycerol, 50 µM ZnCl_2,_ 5 mM beta-mercaptoethanol; 10 mL per gram wet cell pellet) freshly supplemented with 1x PIC (Roche ref. 11873580001), 1x AEBSF (Fisher scientific ref. 10742885), 1x DNaseI (Roche ref. 10104159001), 0.5 mM CaCl_2_, 4 mM MgCl and 500 µg/mL lysozyme (Sigma Aldrich ref. 62971-10G-F). Cells were incubated rotating at 4°C before being sonicated (two seconds off, two seconds on, for a total of 20 seconds at 50% amplitude). Samples were centrifuged to remove cell debris (39,000 x *g*, 30 minutes, 4°C) and the supernatants containing soluble proteins were filtered before being affinity purified on an ÄKTA pure chromatography system using a pre-equilibrated IMAC column (Cytiva ref. 17092102; equilibrated in lysis buffer). The column was washed with 20 CV of 20 mM imidazole and 10 CV of 30 mM imidazole, then eluted in a block with 500 mM imidazole. Pooled fractions containing protein were desalted over a HiPrep 26/10 column (Cytiva ref. 17508701, pre-equilibrated in SEC buffer: 150 mM NaCl, 50 mM Tris pH 7.5, 10% (v/v) glycerol, 50 µM ZnCl_2_, 5 mM beta-mercaptoethanol) to remove imidazole. Then the samples were applied to ion exchange columns (SP and Q attached together, Cytiva ref. 17115201 and 17515601) pre-equilibrated in SEC buffer. The columns were washed separately with 10 CV of SEC buffer and proteins eluted across a 20 CV gradient from 150 mM to 1 M NaCl. Only proteins from SP elutions and washes as well as the SPQ flow-through were kept for later purification while proteins that bound to the Q column were discarded due to DNA contamination. After an additional desalting run over the HiPrep 26/10 column (Cytiva ref. 17508701) for SP elution fractions, samples were purified by gel filtration (Superdex 200 pg, HiLoad 16/600, Cytiva ref. 28989335, pre-equilibrated in SEC buffer) and fractions containing QID-ZFC4 fusion proteins were identified by SDS-PAGE before being pooled, concentrated again and frozen in aliquots at –70°C.

#### SEC-MALS

For the minimal QID peptide, 100 µL of SALL4 was run (at 2.97 mg/mL and 0.97 mg/mL) on a Superdex-75 Increase 10/300 GL size exclusion column pre-equilibrated in buffer containing 25 mM Tris pH 8.0; 250 mM NaCl, 5% (v/v) glycerol and 2.5% (v/v) sucrose at 22°C with a flow rate of 0.9 mL/min. Size-exclusion chromatography (ÄKTA pure, Cytiva) coupled with UV, static light scattering and refractive index (RI) detection (Viscotek SEC-MALS 20 and Viscotek RI Detector VE3580; Malvern Instruments, Malvern, Worcestershire, UK) were used to determine the absolute molecular mass of each protein/complex in solution. Light scattering, RI and A280 nm were analysed by a homo-polymer model (OMNIsec software, v5.02; Malvern Instruments) using the supplied extinction coefficients for ∂A280nm/∂c = 2.03 AU·mL^−1^·mg^−1^, ∂n/∂c = 0.19 mL·g^−1^ for protein and a buffer RI value of 1.34.

For the QID-ZFC4 fusion peptides the experimental procedure was similar, but the running buffer had a different composition (150 mM NaCl, 50 mM Tris pH 7.5, 1% (v/v) glycerol, 50 µM ZnCl_2_, 0.5 mM beta-mercaptoethanol) with a buffer RI value of 1.337. Each fusion protein was run at 2 mg/mL, 1 mg/mL and 0.5 mg/mL, of which the 1 mg/mL runs are shown.

#### Surface plasmon resonance (SPR)

A S-series SA chip was conditioned by three 60 second injections, at 30 µL/min of 1 M NaCl; 50 mM NaOH, and then 180 seconds of running buffer (50 mM Tris, pH 8.0; 150 mM NaCl; 5 µM ZnCl_2_; 1% glycerol; 0.5 mM beta-mercaptoethanol). Biotinylated DNA (see Methods S1; SPR oligos) was then immobilised to 26 RU, at 5 µL/min in running buffer on Flow-cell 2. Flow-cell 1 was “blocked” by a 30 second injection of 1 mM desthiobiotin in running buffer. Single-cycle kinetic runs were performed using a 2-fold dilution series from 25 µM to 1.56 µM of the indicated protein, in running buffer at 25°C with a flow rate of 30 µL/min in running buffer, with a 30 second association and 60 second dissociation phase, over Fc-1 and Fc-2, bulk subtracting the response on Fc-1. Sensorgrams were double referenced subtracted and data fit to a single site, bimolecular kinetic model, with mass transport considerations.

#### Structural modelling

Modelling of the SALL4 QID (G205-W237, mouse SALL4) was performed using the ColabFold interface^37^, which also utilises AlphaFold-Multimer^36^, with default parameters and a defined stoichiometry of 4x molecules. Interaction interface analysis was done using PDBePISA^93^ and by uploading the model ranked first by AlphaFold-Multimer. Figures were then prepared using PyMOL^94^.

### QUANTIFICATION AND STATISTICAL ANALYSIS

#### Automated quantification of AP-stained colonies

An ImageJ plugin (https://doi.org/10.5281/zenodo.16895130) was used to quantify alkaline phosphatase (AP)-stained colonies from scanned images. User-defined parameters, including minimum pixel area and colony circularity threshold, were set to optimize colony detection. Candidate colonies were segmented and identified using the Cellpose^95^ algorithm, providing automated counts and measurements of individual colony areas.

#### Analysis of TBS-causing and neutral SALL1 variants in humans

Disease-causing SALL1 variants (nonsense and frameshift) were downloaded from ClinVar^38^ (https://www.ncbi.nlm.nih.gov/clinvar/) and only considered when they had TBS or SALL1-related disorders associated with it (condition) as well as a “likely pathogenic” or “pathogenic” tag and at least 1/4 review status stars. Publications associated with specific variants are cited in Table S1. The mutational hotspot region was defined in a previous publication^47^. Neutral, non-disease causing SALL1 variants (predicted loss-of-function, pLoF) were obtained from the gnomAD^48^ database (v4.1.0, https://gnomad.broadinstitute.org/), with the following selection criteria: >2 cases found for each specific variant, no flag associated with dubious sequencing quality or possible pathogenicity. All variants were analysed in Microsoft Excel (v16.80) and visualised in GraphPad/Prism (v9).

#### Quantification of foci/nucleoplasm ratios from live-cell imaging data

Quantification of foci/nucleoplasm ratios from live-cell images was performed using a custom-made ImageJ plugin (https://doi.org/10.5281/zenodo.16761300). In brief, the plugin uses the Cellpose^95^ “nucleitorch_3” model to segment nuclei using DNA staining (Hoechst fluorescence channel, mean z-projection) as input. Background subtraction followed by automatic intensity thresholding were then used to segment foci within the nuclei. Intensities and areas of the foci and nucleoplasm were reported for the Hoechst and SALL4-mScarlet3 channels. The total intensities of the nuclei were reported for SALL1-GFP as a measure of the level of transfection in each cell.

#### CUT&RUN analysis

Paired-end sequencing reads were first trimmed to remove adapter sequences and low-quality bases with TrimGalore^96^ (v0.6.7). Trimmed reads were then aligned to the mouse reference genome (mm10) to generate coordinate-sorted alignment files with Bowtie2^97^ (v2.5.1). In parallel, the same reads were mapped to the *E. coli* K12 genome, which served as a spike-in control. The number of *E. coli* reads per library was used to calculate sample-specific normalisation factors. For each condition, biological replicates were merged to produce consolidated alignment files, which were indexed and summarised for alignment statistics. To visualise genome-wide signal, coverage tracks were generated from the merged BAM files. Properly paired fragments shorter than 1 kb were retained, blacklisted regions were excluded, and coverage was calculated across the genome. Tracks were normalised using both RPKM and the spike-in scaling factor, yielding bigWig files that were directly comparable across conditions. Peaks of enrichment were identified with MACS2^98^ (v2.2.7.1) by comparing treatment samples against IgG controls, using paired-end mode and allowing all duplicate fragments to be retained, as appropriate for CUT&RUN data.

Genome-wide binding was assessed by partitioning the mouse genome into non-overlapping 1 kb windows and computing log2-transformed RPKM values from CUT&RUN. For each window, the AT fraction was calculated and windows were ranked by AT content (40–80%). To visualise enrichment relative to AT richness, per-genotype signals were normalised to S4KO (log2 ratio to S4KO) and displayed as smoothed profiles (rolling mean over 5,000 windows, sampled every 2,000). Shaded bands reflect local variability (rolling variance).

Command line arguments and source code for analysis is detailed in Methods S2.

#### RNA-seq analysis

Reads were first aligned to a reference genome of mouse rDNA to remove ribosomal RNA contamination using Bowtie2^97^ (v2.5.1) followed by alignment to mouse reference genome with the STAR aligner^99^ (v2.7.10b). Post-alignment, the featureCounts^100^ tool (v2.0.6) was employed to quantify gene expression levels and differential gene expression analysis was performed using standard DESeq2^101^ (v1.42.0) workflows, with adjusted p-values used to determine statistical significance (expression in mutant lines versus WT). Command line arguments and source code for analysis is detailed in Methods S2.

## Supporting information

Key resources table

Bioinformatic analysis

## DATA AVAILABILITY

Reagents are available upon request.

Raw and processed high-throughput sequencing data generated for this study are available in the GEO database under the accession number GSE306601.

Custom-made ImageJ plug-ins for automated analysis of AP-stained colonies and SALL4 foci-nucleoplasm ratios from live-cell imaging have been deposited on Zenodo (https://doi.org/10.5281/zenodo.16895130 and https://doi.org/10.5281/zenodo.16761300).

## Acknowledgements

This work was supported by funding for the Wellcome Centre for Cell Biology (203149) and the Wellcome Discovery Research Platform for Hidden Cell Biology (226791) and we gratefully acknowledge support from the Light Microscopy and Proteomics cores. We also thank the Edinburgh Protein Production Facility (University of Edinburgh, UK, funded by the School of Biological Sciences) for access to facilities and support. We are grateful to Dr Fiona Rossi at the Flow Cytometry facility at the Institute for Regeneration and Repair (University of Edinburgh, UK) for flow cytometry support and the University of Edinburgh Central Transgenic Core (CTC) for production of mouse lines by blastocyst injection. We thank Dr Brian Hendrich and Prof Ian Chambers for the sharing of cell lines and Dr Marcus Wilson and Dr Hannah Wapenaar for sharing plasmids. We are grateful to Dr Chris Wood for helpful discussions concerning the mode of interaction of SALL proteins. Lastly, we thank Dr. George-Abraham and genetic councillors at Dell Children’s Medical Group Far West for the sharing of patient data for the OS-causing missense mutation in the SALL4 QID as well as GeneDx for establishing contact.

This work was supported by a Wellcome Investigator Award (107930) and an ERC Gen-Epix grant (694295) awarded to A.B.

## Author contributions

Conceptualisation, A.B., R.P., S.G. and K.C.; Methodology, S.G., R.P., K.C., M.W., H.B., J.G., T.Mc., D.K. and C.S.; Software, K.C., T.Mc. and D.K.; Formal Analysis, K.C., T.Mc., D.K. and S.G.; Investigation, S.G., R.P., K.C., M.W., H.B., J.G., T.M., L.G., G.A. and C.S.; Writing – Original Draft, S.G., R.P. and A.B.; Writing – Review & Editing, S.G., K.C., M.W., J.G., H.B., L.G., G.A., C.S., T.Mc., D.K., R.P. and A.B.; Supervision, R.P. and A.B.; Funding acquisition, A.B.

## Supplementary data

**Figure S1.**
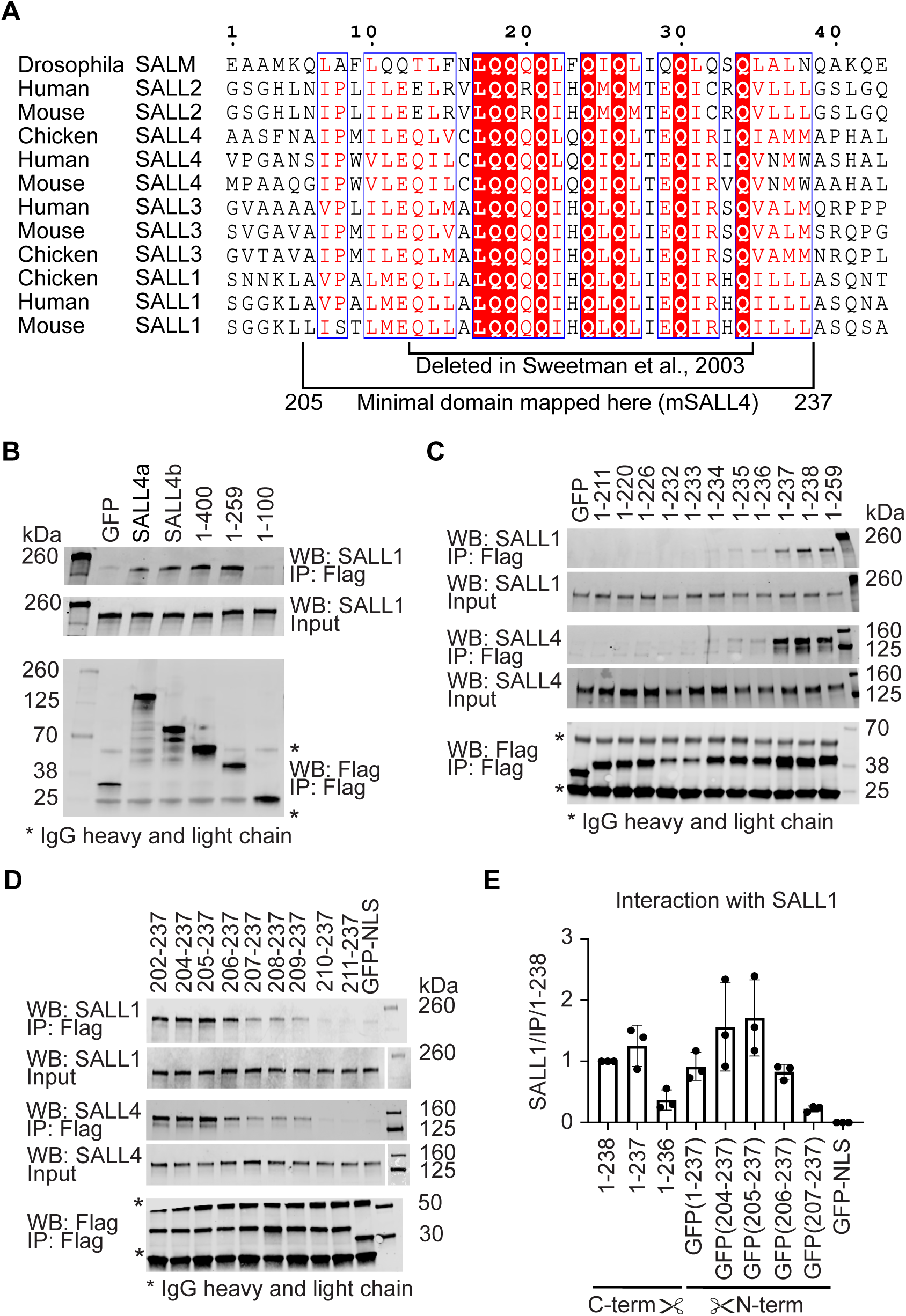
Characterisation of a minimal tetramerisation domain in SALL4, related to Figure 1. (A) Sequence alignment of the Q-rich regions of SALL proteins. Sequences were retrieved from UniProt,^102^ aligned in Clustal Omega^103^ and coloured using ESPript 3.0^104^. White letters on a red background mark identical residues while conservative substitutions are written as red letters. The region shown here corresponds to mouse SALL4 methionine 200 to leucine 242. (B) Western blot analysis of co-IPed endogenous SALL1 from transfections of early C-terminal truncation constructs as well as SALL4a and SALL4b isoforms into E14/T ESCs. All constructs were 3xFlag-tagged and IPed using anti-Flag magnetic beads. Asterisks indicate IgG heavy and light chains of the mouse anti-Flag antibody. (C) Western blot analysis of co-IPed endogenous SALL1 and SALL4 from transfections of the final C-terminal truncation constructs into E14/T ESCs. All constructs were 3xFlag-tagged and IPed using anti-Flag magnetic beads. Asterisks indicate IgG heavy and light chains of the mouse anti-Flag antibody. (D) Western blot analysis of co-IPed endogenous SALL1 and SALL4 from transfections of the final N-terminal truncation constructs into E14/T ESCs. All constructs were 3xFlag-monoGFP-tagged and IPed using anti-Flag magnetic beads. Asterisks indicate IgG heavy and light chains of the mouse anti-Flag antibody. (E) Quantification of the interaction of truncated SALL4 constructs with endogenous SALL1 from co-IP western blot experiments. The data shown in this graph is from three independent replicate experiments (data points) and error bars show the SD.

**Figure S2.**
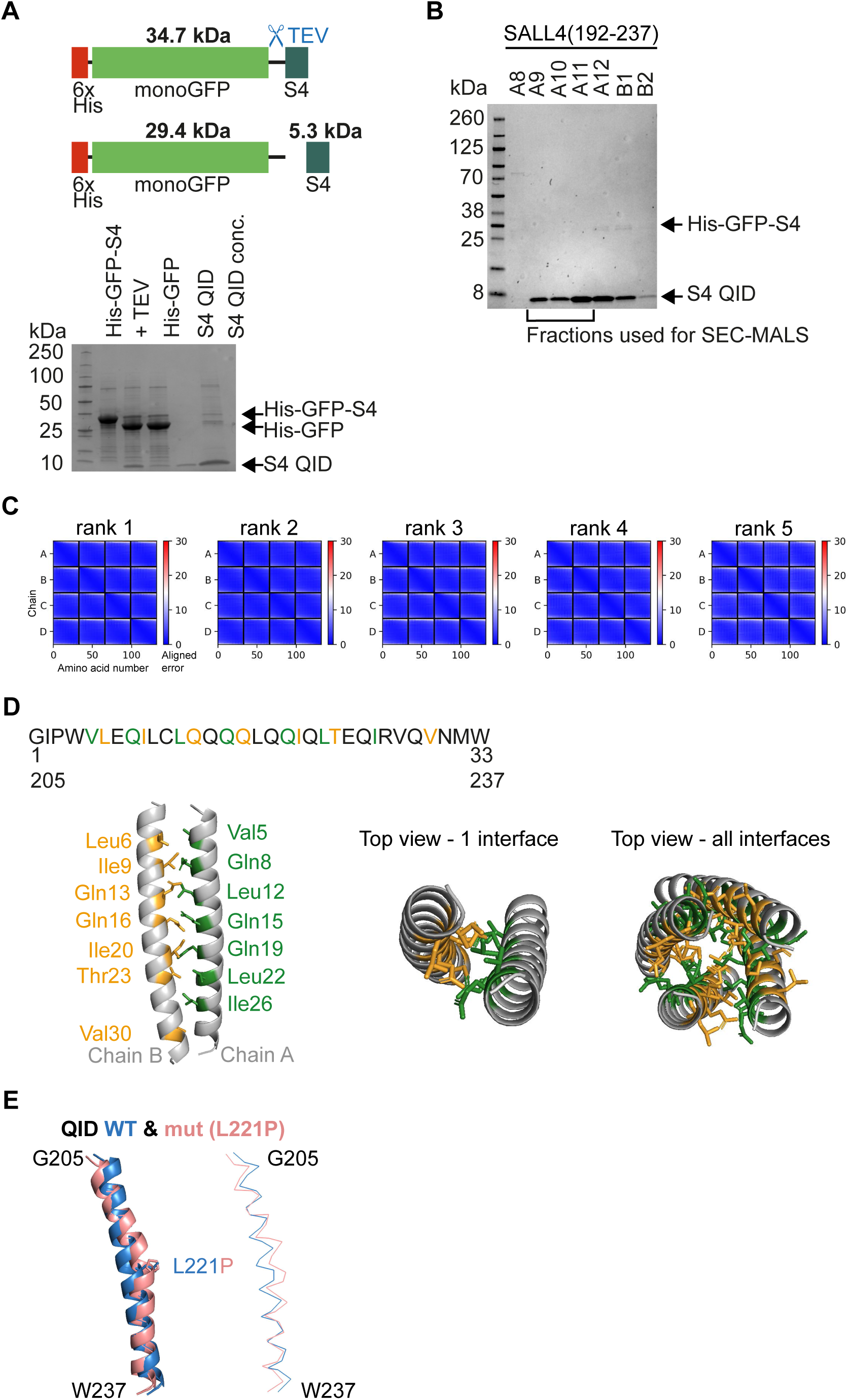
Characterisation of the minimal multimerisation domain in SALL4, related to. Figure 1 (A) Diagram of the recombinant protein that was used for expression in *E. coli* and subsequent purification. Addition of TEV protease removed the 29.4 kDa solubility tag from the 5.3 kDa SALL4 (S4) QID (residues 192-237). Below, a Coomassie staining of a protein gel of the early purification steps is shown with arrows indicating the molecular weight of the different proteins (B) Gel filtration fractions of the SALL4 QID peptide. Arrows indicate the sizes of cleaved SALL4 QID as well as 6xHis-GFP-S4 fusions. The bracket shows which fractions were used for the experiments. (C) Predicted aligned error plots for all five AlphaFold2 models of the SALL4 QID showing a low error rate for the entirety of the region in all models. (D) PDBePISA interface analysis of the rank1 AlphaFold2 model of the QID tetramer.^93^ On top, the sequence of the minimal SALL4 interaction domain (G205-W237) and the numbers corresponding to the AlphaFold2 tetramer model (1-33) are shown. Residues that have a high buried surface area (BSA), meaning that they contribute to the interface, are highlighted in yellow for chain B and green for chain A. Below, the AlphaFold2 model for one interface between chain A (green) and chain B (yellow) is shown from the side and the top, as well as a top view of all the interfaces in the tetramer. (E) Overlay of the rank one AlphaFold2 models of the WT SALL4 QID and the corresponding region with the L221P mutation (QIDmut). WT is shown in blue and L221P in salmon. The line model shows the distortion that is introduced by a proline mutation in the middle of the helix.

**Figure S3.**
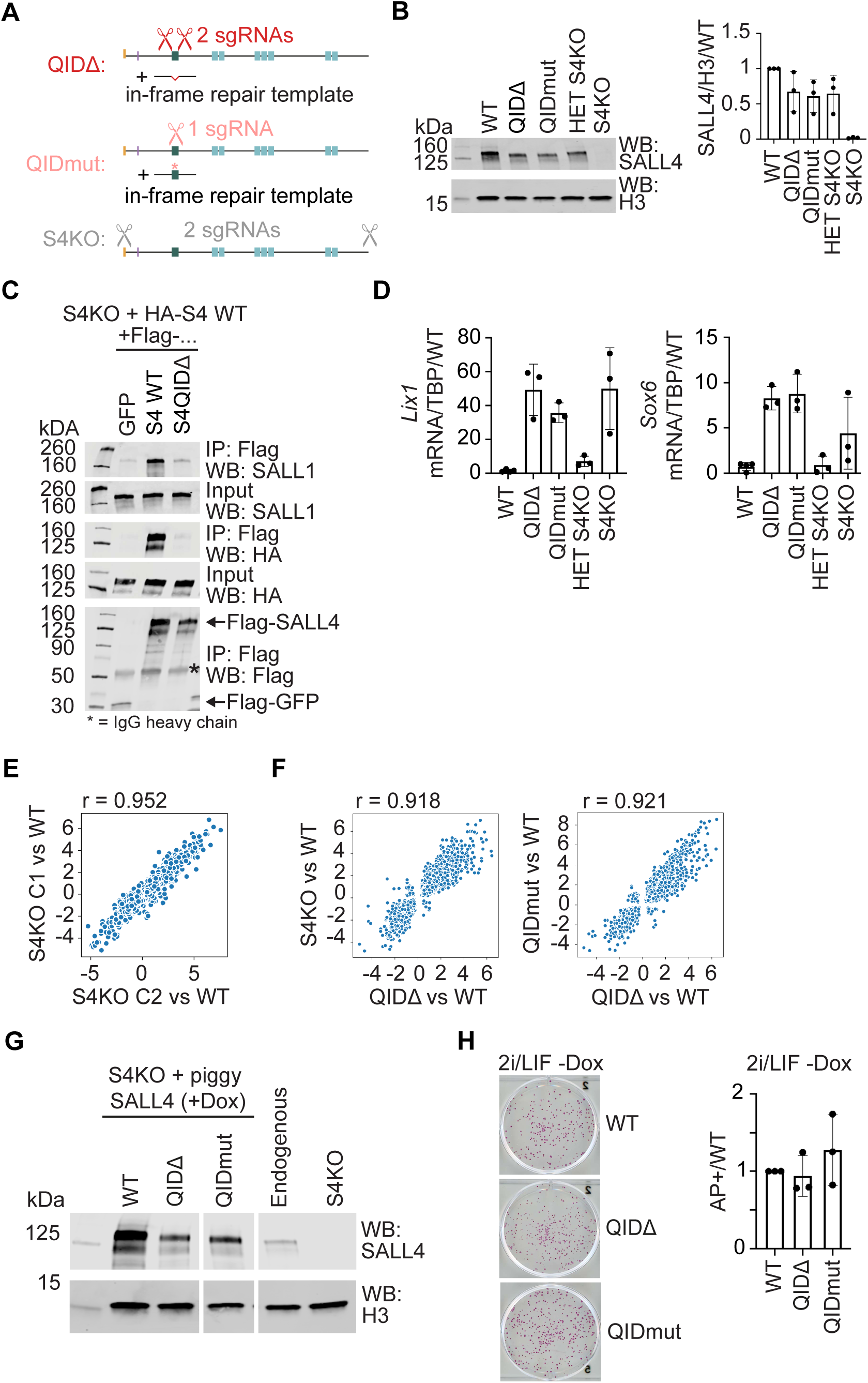
Multimerisation-deficient SALL4 phenocopies a full SALL4 knockout, related to. Figure 2 (A) Diagram depicting the targeting strategies of the endogenous *Sall4* with CRISPR/Cas9 to introduce the deletion of the QID (QIDΔ), the L221P point mutation (QIDmut) and a full *Sall4* KO (S4KO). (B) Western blot analysis of whole cell lysates from the QIDΔ, QIDmut and control cell lines. The quantification on the right shows the mean from three replicate experiments with error bars representing the SD. (C) Western blot from co-IP experiments where 3xFlag-tagged SALL4 versions (WT or QIDΔ) or 3xFlag-GFP were transfected into S4KO ESCs alongside an HA-tagged version of SALL4 full-length. Constructs were IPed with anti-Flag beads and the asterisk indicates the IgG heavy chain of the Flag antibody. (D) RT-qPCR results using cDNA from the different genetically modified cell lines and primer pairs to detect *Lix1* and *Sox6*. All values were normalised to a TBP value from the same sample (internal control) and scaled to WT for comparison. All samples were analysed in technical triplicates to obtain an average and the experiment was repeated three times for biological replicates. The error bars indicate the SD between biological replicates. (E) Plot depicting the correlation of log_2_ fold changes for S4KO C1 and C2. The comparison is done on 3,419 deregulated genes that are shared between each S4KO C1, C2, QIDΔ and QIDmut and the Pearson correlation value (r) is shown. (F) Plots depicting the correlation of log_2_ fold changes of the 2,938 deregulated genes between each genotype in Figure 2D. For each plot the Pearson correlation value (r) is shown. (G) Whole cell lysates of piggyBac cell lines cultured in normal ES cell medium +Dox and control cell lines probed in a western blot against SALL4 and H3. The row labelled endogenous indicates levels in WT E14 ESCs.^77^ (H) Representative wells of stem cell colonies stained by alkaline phosphatase staining^44^ and quantification of all the different piggyBac cell lines that have been cultured in 2i/LIF -Dox medium. For details see Figure 2F.

**Figure S4.**
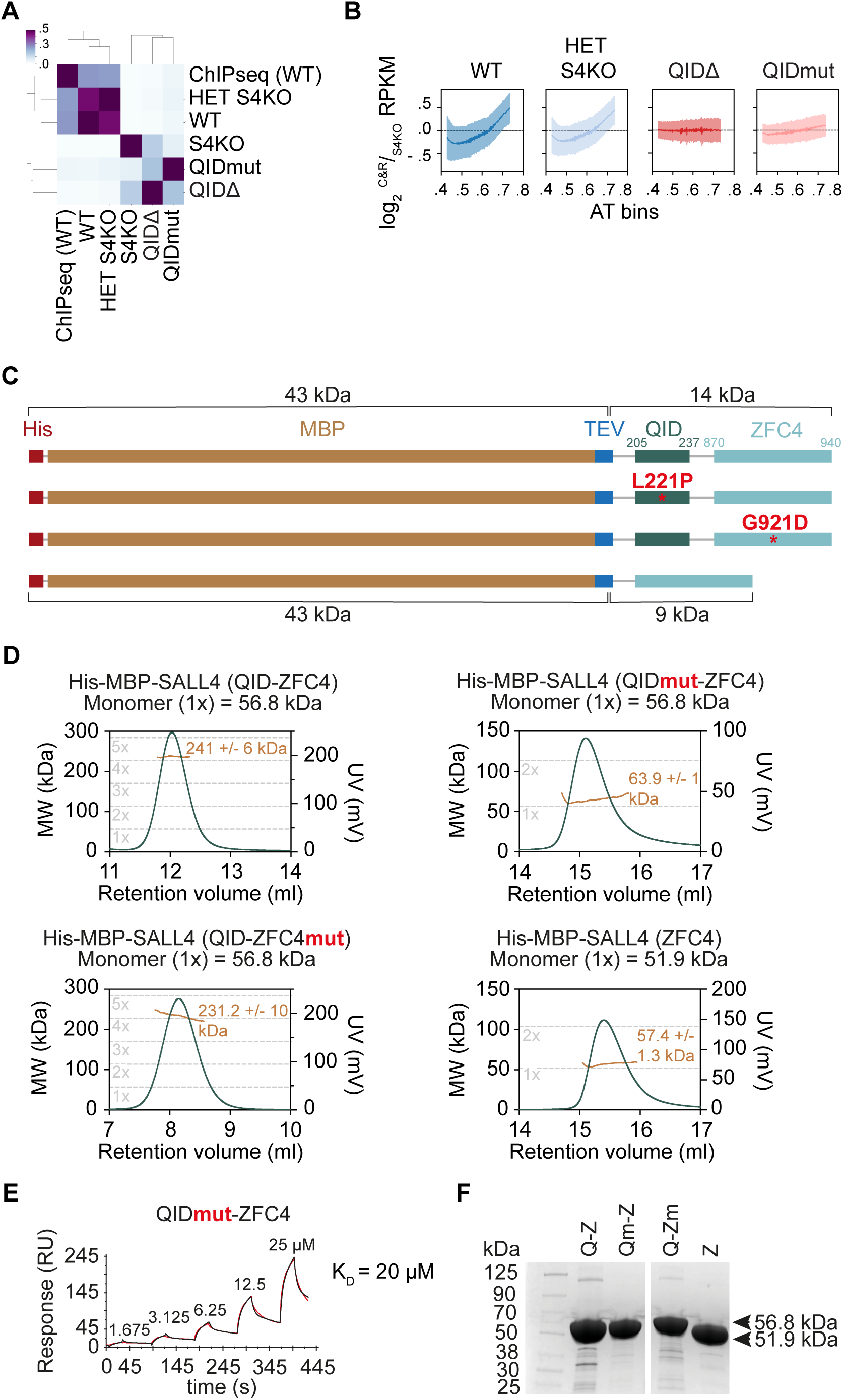
Monomeric SALL4 cannot bind to DNA efficiently, related to. Figure 3 (A) Jaccard overlap of the CUT&RUN peaks found in the different genotypes and SALL4 WT peaks found in a previous study.^8^ The colour scale indicates the Jaccard index from 0 (light blue) to 0.5 (purple), with higher values indicating a stronger overlap between the samples. (B) Log_2_ ratio of the SALL4 CUT&RUN signal (Reads per Kilobase per Million mapped reads (RPKM)) normalised to the S4KO negative control. RPKM were calculated in 1 kb windows and plotted against their AT content in bins ranging from 0.4 (less AT than GC) to 0.8 (more AT than GC). Opaque regions indicate the variance (square root of the SD). (C) Diagram depicting His-MBP-SALL4(QID-ZFC4) fusion constructs that were purified in *E. coli* and their molecular weights. Mutations in functional domains are highlighted with a red *. (D) SEC-MALS analysis of SALL4 (QID-ZFC4) fusion proteins. Green line indicates the protein coming off the SEC while the orange line shows the molecular weight that was measured by the MALS. Grey dashed lines indicate the molecular weights of monomer (1x), dimer (2x), trimer (3x), tetramer (4x) and pentamer (5x) assemblies. (E) SPR analysis of SALL4 QIDmut-ZFC4 fusion protein binding to a DNA probe containing a single ATATTA motif. A K_D_ calculated from a kinetic model is given. (F) Protein gel stained with Coomassie of the purified and concentrated His-MBP-S4(QID-ZFC4) fusion proteins.

**Figure S5.**
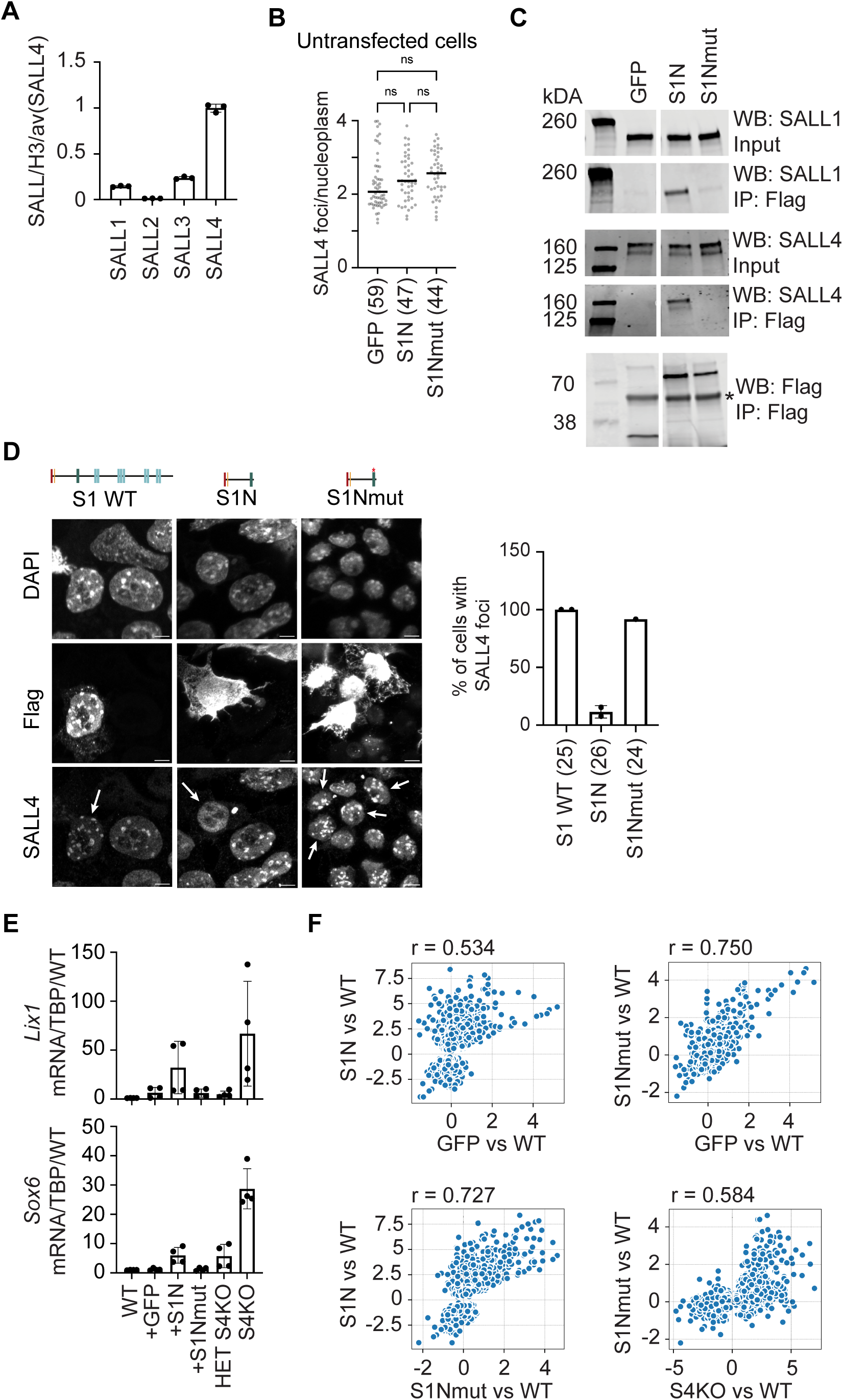
A TBS-causing SALL1 truncation impairs SALL4 sub-nuclear localisation and gene regulation, related to. Figure 5 (A) Nuclear proteome extracts from WT ESCs analysed by mass spectrometry. iBAQ values of the SALL proteins were normalised by the iBAQ value of H3.1 in the same replicate. All values were scaled to the average of SALL4 for comparison and error bars show the SD of the technical triplicates. (B) Quantification of all replicate live-cell imaging experiments. Only cells that were not transfected are shown here with each data point representing a cell and the black line representing the median of the data. Statistical analysis was performed using an ordinary one-way ANOVA with a p-value less than 0.05 being considered statistically significant. (C) Anti-Flag IP of SALL1 truncations or GFP followed by western blot detection of interaction with endogenous SALL1 or SALL4. Note that the endogenous SALL4 is now bigger in size due to the addition of the mScarlet3 tag. The truncated SALL1 constructs were detected with an anti-Flag antibody as the epitope of the SALL1 antibody is at the very C-terminus and thus is lost in this truncation. * Indicates IgG heavy chains. (D) Representative immunofluorescence microscopy pictures of ESCs transfected with GFP, S1N or S1Nmut. White arrows in the SALL4 column indicate transfected cells. These images are from fixed samples and the scale bar represents 5 µm. Quantification of two replicate experiments shown on the right with datapoints representing the percentage of cells with foci from one experiment. Error bars depict the SD while the numbers in brackets show the total number of analysed nuclei. (E) RT-qPCR results using cDNA from WT ESCs transfected with either GFP, S1N or S1Nmut and primer pairs to detect SALL4 target genes *Lix1* and *Sox6*. All values were normalised to a TBP value from the same sample (internal control) and scaled to WT for comparison. They were analysed in technical triplicates to obtain an average and the experiment was repeated four times for biological replicates. The error bars indicate the SD between biological replicates. (F) Plots depicting the correlation of log_2_ fold changes of the 4,317 shared deregulated genes between S1N and S4KO for all other tested samples. For each plot the Pearson correlation coefficient (r) is shown.

**Table S1.** A TBS-causing SALL1 truncation impairs SALL4 sub-nuclear localisation and gene regulation, related to Figure 5. Table showing all indels and nonsense mutations in SALL1 that lead to premature stop codons and the potential loss of functional domains (QID and ZFCs indicated in the table). The minimal QID is based on the mapping experiments in SALL4 and extrapolated to SALL1 by sequence conservation (L224-L257). All variants are from the ClinVar^38^ database or previous publications where indicated, which means that they are associated with a disease-phenotype (TBS or SALL1-related disorder). The mutations indicate the direct base changes found in human SALL1 (ENST00000440970.6) and the corresponding protein changes are shown next to it. * indicate stop codons while fs stands for frameshifts with the number after the stop codon showing the length of the missense tail. The cases column indicates the number of patients that have been found with this variant.

| **Mutation** | **Protein consequence** | **QID retained?** | **ZFCs lost?** | **Cases** | **Reference** | **ClinVar<sup>38</sup> identifier** |
| --- | --- | --- | --- | --- | --- | --- |
| c. 252del | p. F85S fs*14 | No | All | 1 | None | VCV003236751.1 |
| c. 269dup | p. P91S fs*2 | No | All | 1 | None | VCV001048611.1 |
| c. 313del | p. T105Q fs*76 | No | All | 1 | Botzenhart, 2007 <sup>47</sup> | None |
| c. 350del | p. N117T fs*66 | No | All | 1 | None | VCV003725389.1 |
| c. 388_391delins | p. P130F fs*61 | No | All | 1 | None | VCV001179917.2 |
| c. 420del | p. S141A fs*42 | No | All | 1 | Kohlhase, 1999 <sup>28</sup> | VCV000934902.9 |
| c. 469_512dup | p. I172A fs*26 | No | All | 1 | None | VCV002103715.3 |
| c. 601C>T | p. Q201* | No | All | 2 | None | VCV001679304.5 |
| c. 691del | p. E231N fs*4 | No | All | 1 | None | VCV001353868.5 |
| c. 712C>T | p. Q238* | No | All | 1 | None | VCV000562380.7 |
| c. 724del | p. H242T fs*5 | No | All | 1 | None | VCV003236775.1 |
| c. 750dup | p. R251S fs*19 | No | All | 1 | None | VCV000947631.9 |
| c. 764del | p. L255 fs | No | All | 2 | Botzenhart, 2005 <sup>105</sup> | None |
| c. 778C>T | p. Q260* | Yes | All | 2 | Botzenhart, 2005 <sup>105</sup> | None |
| c. 792_793del | p. L264 fs* | Yes | All | 5 | Surka, 2001 <sup>106</sup> | VCV000007433.1 |
| c. 814C>T | p. Q272* | Yes | All | 1 | Botzenhart, 2007 <sup>47</sup> | VCV000488838.9 |
| c. 817del | p. G273 fs* | Yes | All | 1 | Salerno, 2000 <sup>107</sup> | None |
| c. 826C>T | p. R276* | Yes | All | 10 | Kohlhase, 1999 <sup>28</sup> | VCV000007428.21 |
| c. 840del | p. L282 fs* | Yes | All | 1 | Blanck, 2000 <sup>108</sup> | None |
| c. 866T>A | p. L289* | Yes | All | 1 | None | VCV000528879.8 |
| c. 870dup | p. Q291S fs*21 | Yes | All | 1 | None | VCV001075840.6 |
| c. 870_871dup | p. Q291L fs*24 |  |  | 1 |  | VCV000279887.9 |
| c. 871C>T | p. Q291* |  |  | 2 |  | VCV000936315.10 |
| c. 874C>T | p. Q292* | Yes | All | 1 | Lin, 2016 <sup>109</sup> | VCV002137807.5 |
| c. 878_887del | p. L293Q fs*18 | Yes | All | 1 | None | VCV002031602.5 |
| c. 881_893dup | p. L299S fs*17 | Yes | All | 1 | None | VCV001070663.9 |
| c. 901C>T | p. Q301* | Yes | All | 1 | Botzenhart, 2007 <sup>47</sup> | None |
| c. 928del | p. I310L fs*4 | Yes | All | 1 | None | VCV003728154.1 |
| c. 940A>T | p. K314* | Yes | All | 1 | Botzenhart, 2005 <sup>105</sup> | None |
| c. 952_953del | p. P318N fs*36 | Yes | All | 1 | None | VCV003764693.1 |
| c. 953del | p. P318Q fs*24 |  |  | 1 |  | VCV003062088.1 |
| c. 958C>T | p. Q320* | Yes | All | 1 | None | VCV000528880.8 |
| c. 967C>T | p. Q323* | Yes | All | 5 | Albrecht, 2004 <sup>110</sup> | VCV000007434.9 |
| c. 995del | p. P332H fs*10 | Yes | All | 1 | Botzenhart, 2007 <sup>47</sup>;<br>Furniss, 2007 <sup>111</sup> | VCV000007436.1 |
| c. 1027dup | p. I343N fs*12 | Yes | All | 1 | None | VCV001071229.8 |
| c. 1028_1029del | p. I343 fs*9 |  |  |  | Botzenhart, 2005 <sup>105</sup> | None |
| c. 1028dup | p. L344I fs*11 | Yes | All | 1 | None | VCV001069953.6 |
| c. 1045dup | p. T349N fs*5 | Yes | All | 1 | Botzenhart, 2007 <sup>47</sup> | None |
| c. 1047dup | p. T350H fs*4 | Yes | All | 2 | Botzenhart, 2007 <sup>47</sup> | None |
| c. 1061dup | p. E354 fs*35 | Yes | All | 5 | Botzenhart, 2007 <sup>47</sup> | None |
| c. 1084_1085del | p. A362L fs*26 | Yes | All | 1 | Botzenhart, 2007 <sup>47</sup> | None |
| c. 1108_1109del | p. V370L fs*19 | Yes | All | 1 | None | VCV000426114.3 |
| c. 1115C>G,A | p. S372* | Yes | All | 3 | Kohlhase, 1998 <sup>112</sup>;<br>1999 <sup>28</sup> | VCV000007429.2<br>VCV000007427.1 |
| c. 1119_1197del | p. S373 fs*17 | Yes | All | 1 | Botzenhart, 2007 <sup>47</sup> | None |
| c. 1124del | p. S375Y fs*5 | Yes | All | 1 | Botzenhart, 2005 <sup>105</sup> | None |
| c. 1134del | p. F378 fs* | Yes | All | 1 | Botzenhart, 2007 <sup>47</sup> | None |
| c. 1148del | p. L383Y fs*2 | Yes | All | 1 | None | VCV002687773.1 |
| c. 1145_1146ins | p. L383I fs*2 |  |  | 1 | Botzenhart, 2005 <sup>105</sup>;<br>Kohlhase, 1999 <sup>28</sup> | None |
| c. 1146del | p. L383Y fs*2 |  |  | 1 |  |  |
| c. 1174_1175del | p. L392T fs*6 | Yes | All | 2 | Botzenhart, 2007 <sup>47</sup> | VCV003899973.1 |
| c. 1183C>T | p. Q395* | Yes | All | 1 | None | VCV002694137.2 |
| c. 1199C>A | p. S400* | Yes | All | 1 | None | VCV003648236.1 |
| c. 1200_1206del | p. V401P fs*13 | Yes | All | 2 | Kohlhase, 1999 <sup>28</sup> | None |
| c. 1214dup | p. L406F fs*32 | Yes | All | 1 | None | VCV000459257.6 |
| c. 1228G>T | p. G410* | Yes | All | 2 | Botzenhart, 2007 <sup>47</sup> | VCV001705680.1 |
| c. 1256T>A | p. L419* | Yes | All | 3 | Kosaki, 2007 <sup>113</sup>; | VCV000007435.1 |
| c. 1255del | p. L419C fs*73 |  |  | 1 | Devriendt, 2002 <sup>114</sup> | None |
| c. 1263del | p. L422W fs*70 | Yes | All | 1 | Botzenhart, 2005 <sup>105</sup> | None |
| c. 1270del | p. Q424S fs*68 | Yes | All | 3 | Kohlhase, 1998 <sup>112</sup> | VCV000007426.1 |
| c. 1273del | p. Q425K fs*67 | Yes | All | 2 | Botzenhart, 2007 <sup>47</sup> | None |
| c. 1277_1278del | p. R426K fs*10 | Yes | All | 1 | Kohlhase, 1999 <sup>28</sup> | VCV000007430.1 |
| c. 1291_1300del | p. P431L fs*58 | Yes | All | 1 | Kohlhase, 1999 <sup>28</sup> | None |
| c. 1309G>T | p. E437* | Yes | All | 1 | None | VCV002703340.2 |
| c. 1321dup | p. T441N fs*3 | Yes | All | 1 | Botzenhart, 2005 <sup>105</sup> | None |
| c. 1324del | p. S442P fs*51 | Yes | All | 1 | None | VCV000648831.8 |
| c. 1327del | p. D443M fs*49 | Yes | All | 2 | Botzenhart, 2005 <sup>105</sup> | None |
| c. 1347_1348del | p. H449Q fs*11 | Yes | All | 1 | Marlin 1999 <sup>115</sup> | VCV000007431.1 |
| c. 1363_1369delins | p. A455 fs* | Yes | All | 1 | None | VCV001679359.3 |
| c. 1365_1366ins4 | p. K456A fs*6 | Yes | All | 1 | None | VCV000983477.2 |
| c. 1374del | p. F458L fs*34 | Yes | All | 1 | Botzenhart, 2005 <sup>105</sup> | None |
| c. 1404dup | p. R469A fs*46 | Yes | All | 3 | Botzenhart, 2005 <sup>105</sup> | None |
| c. 1411_1412ins | p. H471Q fs*44 | Yes | All | 2 | Marlin 1999 <sup>115</sup> | None |
| c. 1417G>T | p. G473* | Yes | All | 1 | None | VCV002005719.3 |
| c. 1415_1425del | p. G473I fs*38 |  |  | 1 | Botzenhart, 2007 <sup>47</sup> | None |
| c. 1423_1424del | p. R475A fs*40 | Yes | All | 1 | None | VCV001069244.8 |
| c. 1479_1480ins | p. V494R fs*15 | Yes | All | 2 | Marlin 1999 <sup>115</sup> | None |
| c. 1487_2048del | p. Q497R fs*11 | Yes | All | 5 | Marlin 1999 <sup>115</sup> | None |
| c. 1503_1504del | p. K502I fs*12 | Yes | ZFC2-4 | 1 | Botzenhart, 2007 <sup>47</sup> | None |
| c. 1509C>A | p. Y503* | Yes | ZFC2-4 | 2 | Blanck, 2000 <sup>108</sup> | None |
| c. 1516_1517dup | p. Q507S fs*2 | Yes | ZFC2-4 | 2 | Botzenhart, 2007 <sup>47</sup> | None |
| c. 1543G>T | p. E515* | Yes | ZFC2-4 | 3 | Botzenhart, 2005 <sup>105</sup> | None |
| c. 1565del | p. T522R fs*20 | Yes | ZFC2-4 | 4 | Marlin 1999 <sup>115</sup> | None |
| c. 1606A>T | p. L536* | Yes | ZFC2-4 | 1 | None | VCV003254898.1 |
| c. 1819del | p. L608S fs*34 | Yes | ZFC2-4 | 3 | Engels, 2000 <sup>116</sup> | VCV000007432.1 |
| c. 1873G>T | p. E625* | Yes | ZFC2-4 | 1 | None | VCV000973255.5 |
| c. 1966_1975del | p. T656P fs*36 | Yes | ZFC2-4 | 1 | Marlin 1999 <sup>115</sup> | None |
| c. 2129del | p. I710T fs*11 | Yes | ZFC2-4 | 1 | None | VCV001700011.1 |
| c. 2225_2226del | p. A742V fs*8 | Yes | ZFC2-4 | 1 | None | VCV002630177.1 |
| c. 2256del | p. Y753T fs*69 | Yes | ZFC2-4 | 1 | Botzenhart, 2007 <sup>47</sup> | VCV000459258.8 |
| c. 2283_2284del | p. L762Q fs*41 | Yes | ZFC2-4 | 1 | None | VCV002681784.2 |
| c. 2325_2331dup | p. A778H fs*29 | Yes | ZFC2-4 | 1 | None | VCV002837510.2 |
| c. 2356del | p. R786E fs*36 | Yes | ZFC2-4 | 1 | None | VCV000528881.2 |
| c. 2390del | p. P797Q fs*24 | Yes | ZFC3+4 | 1 | None | VCV003765528.1 |
| c. 2686_2689dup | p. D896S fs*1 | Yes | ZFC3+4 | 1 | None | VCV000807481.3 |
| c. 2779C>T | p. Q927* | Yes | ZFC3+4 | 1 | Walter, 2006 <sup>117</sup> | None |
| c. 3005_3008del | p. A1002V fs*42 | Yes | ZFC3+4 | 1 | None | VCV000663219.1 |
| c. 3019_3032dup | p. C1012A fs*37 | Yes | ZFC3+4 | 2 | Botzenhart, 2007 <sup>47</sup> | VCV003899465.1 |
| c. 3128_3129del | p. N1043I fs*19 | Yes | ZFC3+4 | 1 | None | VCV002828754.2 |
| c. 3160C>T | p. R1054* | Yes | ZFC4 | 3 | Vodopiutz, 2013 <sup>118</sup> | VCV000219151.9 |
| c. 3249_3255del | p. L1084R fs*117 | Yes | ZFC4 | 3 | Botzenhart, 2007 <sup>47</sup> | None |
| c. 3381del | p. R1128G fs*75 | Yes | ZFC4 | 1 | None | VCV001333617.1 |
| c. 3414_3415del | p. C1139W fs*35 | Yes | ZFC4 | 4<br>3 | Botzenhart, 2007 <sup>47</sup>;<br>Furniss, 2007 <sup>111</sup> | VCV000418466.17 |
| c. 3856C>T | p. Q1286* | Yes | None | 1 | None | VCV003235206.1 |

