## Supplementary material for "Tetramerisation governs SALL transcription factor function in development and disease": Bioinformatic analysis

### Bioinformatic analysis – command line arguments

#### CUT&RUN analysis

Sequencing adapters and low-quality bases were removed from paired-end FASTQ files before downstream analysis.

```
trim_galore --basename <SAMPLE> --paired --gzip -o trimmed
<FASTQ_R1> <FASTQ_R2>
```

Trimmed reads were aligned to the mouse reference genome (mm10), and the output was saved as a coordinate-sorted BAM file.

```
bowtie2 -x indexes/mm10 --end-to-end --very-sensitive --no-
unal --no-mixed –no-discordant -1 trimmed/<SAMPLE>_val_1.fq.gz
-2 trimmed/<SAMPLE>_val_2.fq.gz | samtools sort -o
bowtie2/<SAMPLE>.bam -
```

In parallel, the same trimmed reads were also aligned to the *E. coli* K12 genome to measure spike-in DNA and derive a normalisation factor.

```
bowtie2 -x indexes/eschColi_K12 --end-to-end --very-sensitive
--no-unal --no-mixed --no-discordant -1
trimmed/<SAMPLE>_val_1.fq.gz -2 trimmed/<SAMPLE>_val_2.fq.gz |
samtools sort -o spikein/<SAMPLE>.bam -
```

For each condition, replicate BAM files were merged into a single dataset, indexed, and summarised.

```
samtools merge merged/<NAME>.bam <REP1>.bam <REP2>.bam
<REP3>.bam <REP4>.bam <REP5>.bam <REP6>.bam && \

samtools index merged/<NAME>.bam && \

samtools flagstat merged/<NAME>.bam > merged/<NAME>.flagstat
```

The number of *E. coli* reads in each merged library was counted to calculate a sample-specific scaling factor. To visualise CUT&RUN signal across the genome, BAM files were converted into bigWig tracks. Reads were extended, blacklisted regions were excluded, and the coverage was normalised using both RPKM and the spike-in–derived scaling factor. This produced genome-wide signal tracks that were directly comparable across samples.

```
bamCoverage -b bowtie2/<SAMPLE>.bam -bs 10 -e -bl
resources/mm10-blacklist.v2.bed --normalizeUsing RPKM --
smoothLength 30 --scaleFactor <SPIKEIN_FACTOR> --
effectiveGenomeSize 2652783500 --centerReads --samFlagInclude
2 -of bigwig -o scaled_rpkm_bedgraphs/<SAMPLE>.bw
```

Enriched regions of CUT&RUN signal (peaks) were identified by comparing treatment samples against IgG controls.

```
macs2 callpeak -t bowtie2/<SAMPLE>.bam -c
bowtie2/<IGG_SAMPLE>.bam –call-summits -f BAMPE -g mm --keep-
dup all -n <SAMPLE> --outdir macs2_control
```

#### Base composition-dependent genome-wide enrichment of SALL4 CUT&RUN signal

To determine whether SALL4 binding is influenced by base composition, the M. musculus genome was fragmented into non-overlapping 1 kb windows. For each window, the AT fraction was calculated from the reference sequence and the CUT&RUN signal, expressed as log2 RPKM values pre-normalised to S4KO, was quantified for each genotype. Windows were then rank-ordered by their AT fraction to generate an ordered series suitable for trend analysis.

In order to smooth the relationship between AT content and CUT&RUN signal, we applied a rolling mean across 5,000 consecutive windows. For each rolling window, the average CUT&RUN signal was calculated alongside the variance, which provides a measure of local variability around the mean. To reduce redundancy between highly overlapping windows, every 2,000th rolling window was retained for plotting. This procedure yields a smaller set of representative bins, each with an associated mean value and variance, while still capturing the overall trend of the data.

The approach is illustrated below with a toy example in which a rolling mean is calculated across five consecutive elements. Each step shifts the window forward by one element, generating overlapping groups for which the mean and variance are computed. To avoid redundancy, only every third group is retained to plot the final smoothed curve with a shaded variance band.

```
[a b c d e]
  [b c d e f]
    [c d e f g]
      [d e f g h]
        [e f g h i]
          [f g h i j]
            [g h i j k]
              [h i j k l]
                [i j k l m]


    [c d e f g]
          [f g h i j]
                [i j k l m]
```

The same principle was applied to the genomic data using a window size of 5,000 and a step of 2,000. The resulting profiles plot AT fraction on the x-axis against log2 RPKM (relative to S4KO) on the y-axis, with lines representing the smoothed mean CUT&RUN signal and shaded regions indicating variance. This enabled direct comparison of enrichment patterns across genotypes, revealing the dependence of SALL4 binding on underlying base composition. The analysis was implemented in Python and the pseudo code is below:

```
rolling_df = (pd.concat(dfs, axis=1).sort_values(by="AT")
    .rolling(window=5000)
    .mean()[::2000].dropna())

rolling_var_df = (pd.concat(concat,
axis=1).sort_values(by="AT")
    .rolling(window=5000)
    .var()[::2000].dropna())
```

#### RNA-seq analysis

Reads mapping to ribosomal RNA were discarded to avoid bias from highly abundant transcripts. Only unmapped reads were retained for downstream analysis.

```
bowtie2 --no-unal --un-conc-gz
results/rRNA_discarded/<SAMPLE>_%.fastq.gz --very-fast --no-
mixed --no-discordant -x resources/genomes/rRNA -1
fastq/<SAMPLE>_R1_001.fastq.gz -2
fastq/<SAMPLE>_R2_001.fastq.gz > /dev/null
```

The rRNA-filtered reads were aligned to the concatenated reference genome with STAR, which produced an unsorted BAM file along with transcriptome alignments, read counts, and unmapped reads.

```
STAR --genomeDir resources/genomes/concatenated  --readFilesIn
results/rRNA_discarded/<SAMPLE>_1.fastq.gz
results/rRNA_discarded/<SAMPLE>_2.fastq.gz --readFilesCommand
zcat --outSAMtype BAM Unsorted --quantMode TranscriptomeSAM
GeneCounts --twopassMode Basic --outReadsUnmapped Fastx --
outFileNamePrefix results/star/concatenated/<SAMPLE>/
```

Reads that remained unmapped after STAR were re-aligned with Bowtie2 in sensitive local mode to maximise recovery. The resulting alignments were stored in CRAM format.

```
bowtie2 --no-unal --un-conc-gz
results/unmapped/concatenated/<SAMPLE>_%.fastq.gz --sensitive-
local --no-mixed --no-discordant -x
resources/genomes/concatenated -1
results/unmapped/concatenated/<SAMPLE>.1.fastq.gz -2
results/unmapped/concatenated/<SAMPLE>.2.fastq.gz | samtools
sort --reference resources/genomes/concatenated.fa -o
results/sorted/backup/<SAMPLE>_bowtie2.cram -
```

STAR and Bowtie2 alignments were merged into a single CRAM file per sample to obtain complete gene coverage.

```
samtools cat results/sorted/backup/<SAMPLE>.cram
results/sorted/backup/<SAMPLE>_bowtie2.cram | samtools view -h
-F 256 -T resources/genomes/concatenated.fa | samtools sort --
reference resources/genomes/concatenated.fa -o
results/sorted/merged/<SAMPLE>.cram -
```

Read pairs mapping to gene bodies (including introns) were counted using featureCounts. Counts were summarised by gene IDs across all replicates.

```
featureCounts -a results/deduplicated/<GENOME>/genes.saf -F
SAF -o
results/deduplicated/<GENOME>/featureCounts.gene_id.reverse.tx
t -s 2 -p --countReadPairs -B -C -t gene_id -g gene_id
results/deduplicated/<GENOME>/*.bam
```

Gene-level counts were imported into DESeq2. A design formula including both batch and treatment effects was used, and fold changes were estimated with shrinkage for stability.

```
dds <- DESeqDataSetFromMatrix(countData = counts, colData =
design, design = ~ jointly_handled + treatment)
```

```
res_shrink <- lfcShrink(dds, coef = "treatment_vs_wt", type =
"normal")
```
